# Aligned collagen fibers drive distinct traction force signatures to regulate contact guidance

**DOI:** 10.1101/2025.04.22.650031

**Authors:** Gopal Niraula, Azarnoosh Foroozandehfar, Fred Rogers Namanda, Ian Christopher Schneider

## Abstract

Cellular forces on isotropically deposited extracellular matrix (ECM) have been measured extensively. However, *in vivo*, cells exert traction force on collagen fiber networks within ECM. Often times collagen fibers are aligned as in cancer, fibrosis and during wound healing. How forces are transmitted on aligned collagen fibers and how the cytoskeleton regulates this is unknown. Here, we develop a dual-traction force microscopy (d-TFM) approach that includes collagen fibers attached to flexible substrates with fiduciary markers on both the collagen fibers and underlying flexible substrates. This allows for the measurement of traction forces on collagen fibers in the plane of the cell, ensuring a physiologically relevant environment and accurately quantifying traction force. We find that the elastic modulus of the substrate determines the steady-state traction stress exerted by spreading cells on aligned collagen fibers, but does not affect traction force kinetics. Furthermore, collagen fiber networks on the same elastic modulus as isotropically adsorbed collagen result in higher traction stresses. Formins and Arp2/3 modulate traction stress differently, where formins affect traction stress magnitude, while Arp2/3 affects traction stress kinetics. Interestingly, we found that there is a positive correlation between traction force, migration speed and directionality on aligned collagen fibers. However, traction force does not seem to correlate with speed and directionality across distinct collagen organizational structures and across cell lines. These findings underscore the complex interplay between the mechanics of collagen fiber networks, cytoskeletal regulators and cellular traction forces, providing insights into how cells navigate complex fiber networks during migration.

**Graphical Abstract:** 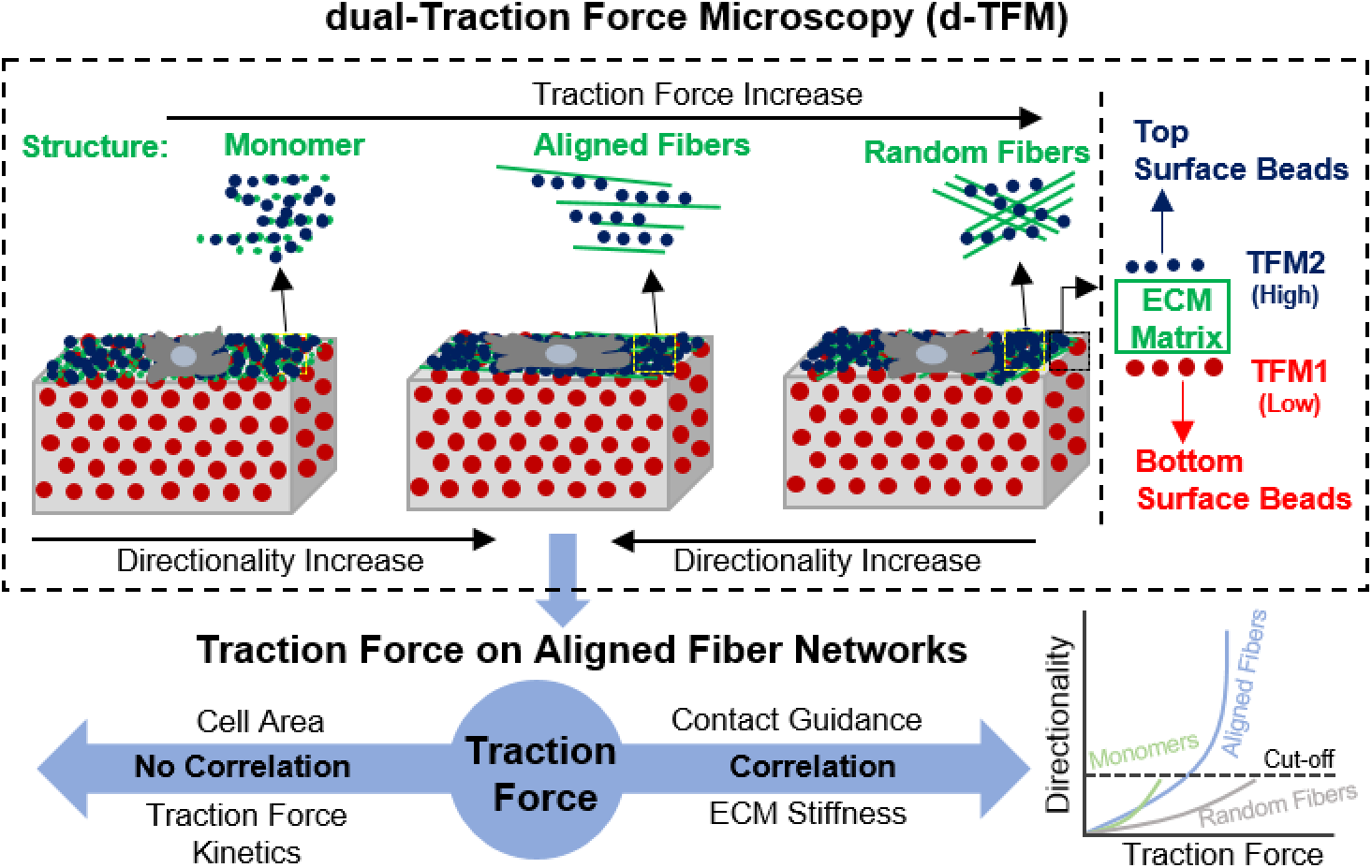

## 1. Introduction

Mechanical force is critical in sensing external environments and transmitting biophysical cues to the cytoskeleton. Biophysical cues control cell-level functions such as adhesion and migration as well as tissue-level functions such as creating specific tissue architecture, encouraging cancer metastasis or developing fibrosis ^1–4^. Cells are typically found within complex collagen fiber networks *in vivo* ^5–8^. The density and architecture of these fiber networks vary among tissues. For example, while normal tissue (connective tissue within skin) contains poorly organized and randomly distributed fibers, metastatic tissue (aggressive breast cancers) contains tightly packed and aligned fibers leading to a directional migration strategy called contact guidance ^9,10^. Cells migrate through these distinct fiber networks by dynamically altering their morphology and attaching to and remodeling the extracellular matrix (ECM). Additionally, cells can change the strategy of their migration, switching between mesenchymal, epithelial and amoeboid migrations modes ^11–14^. A cells ability to adopt a particular morphology, generate and transmit mechanical force and migrate directionally are intimately tied to the structure as well as the stiffness of the ECM. Several studies on isotropic substrates have reported a biphasic relationship between migration speed and stiffness in various types of ECM and cell lines ^15–18^, but how stiffness regulates cell migration on aligned collagen fibers is less well-studied ^19^. Whether this biphasic correlation applies universally across different ECM structures and cell types is yet to be determined. Furthermore, how the cytoskeletal components act to control cell morphology, traction generation and directional migration on fiber networks is not known. These are important questions given that aligned collagen fibers drive so many physiological and pathological processes. Thus, understanding force transmission in aligned and random fiber networks is essential for designing biomaterials, improving regenerative medicine strategies and understanding pathological conditions like cancer invasion and fibrosis.

The assembly and activation of intracellular proteins connecting the actomyosin cytoskeleton to the ECM is critical for understanding force transmission. Focal adhesions (FAs) connected to the F-actin network ^20,21^ transmit force depending upon shape and size as well as cell type-specific levels contractility ^19,22,23^. Force transmitted via FAs and actomyosin contractility are the key factors in controlling cell directionality during contact guidance ^23^. The myosin contractile force further controls migration speed in response to ECM stiffness ^19,24^. Contractile force on fiber networks is enhanced when cells can recruit nearby fibers, even if the substrate is soft ^25^. In addition to FAs and myosin activity, the structure of the F-actin network is another important modulator of cellular force. Cells assemble both linear and branched F-actin networks generating contractile and protrusive force in a controlled fashion, enabling the cell to migrate in response to structural cues. Formins, promote the formation of linear actin filaments, which are essential for creating stress fibers and exerting contractile forces that drive cell body movement and stabilize cell adhesion. Arp2/3 on the other hand is responsible for forming branched actin networks, which are crucial for generating the pushing forces needed for protrusions at the leading edge of migrating cells. While some studies have explored the role of formins and Arp2/3 in contact guidance, these have only been on ridges or microcontact printed lines ^26,27^. It is not understood how formins or Arp2/3 regulate force transmission on aligned collagen fibers. This is important because formins and Arp2/3 likely have different roles during contact guidance. Formins reinforce directed migration by generating F-actin alignment along aligned fibers, whereas Arp2/3 allows the cell to probe the environment in many different directions and make turns. Consequently, understanding how formins and Arp2/3 regulate the force transmission on aligned collagen fibers is essential for revealing fundamental mechanisms of contact guidance.

Traction forces have been measured using traction force microscopy (TFM) ^28,29^, micro-pillars ^30^, and DNA-based force probes ^31^. Traction force microscopy (TFM) has been a tremendous tool for studying the mechanical force exerted by cells on the ECM ^23^. The well-established conventional TFM technology uses isotropically adsorbed ECM molecules on a flexible substrates like polyacrylamide (PAA) gels as the cell adhesive substrate and fluorescence beads as fiduciary markers (FMs) embedded into the PAA gel of desired stiffness ^23,32^. In this approach, cells apply mechanical force and deform the substrate, displacing the beads embedded within the PAA gel. However, two problems exist. First, force transmission on an anisotropic fiber substrate is likely different than on a substrate with isotropically adsorbed ECM monomers. Since cells encounter both aligned and randomly oriented collagen fibers *in vivo*, conventional TFM using collagen monomers does not capture the force transmission characteristics that are specific to interactions between cells and collagen fibers. In addition, conventional TFM assumes beads are at the surface. What is often ignored is force dissipation within PAA gel as a function of depth. Beads far below the surface are unable to measure traction force precisely, which is a particular problem for soft substrates ^33,34^. This results in lower bead deformations at lower depths and an underestimate of the traction force ^32^. In order to both assess physiological traction forces exerted on fibers and to eliminate underestimates of traction force generation, it is important to develop a method to measure cellular traction force directly on aligned collagen fiber networks.

Previously, our lab developed a unique approach of using mica to epitaxially grow various collagen fiber networks and transfer these fibers to substrates for studying cell migration ^35–37^. In this paper, we leverage these approaches to create a dual traction force microscopy (d-TFM) system. This creates aligned or random collagen fiber networks that can be transferred onto flexible substrates with embedded beads that act as FMs. In addition, we can attach beads directly to the fibers creating FMs and enabling traction force measurement on fiber networks. We measure traction stresses over time in spreading cells. We assess the traction stress kinetics and magnitudes on substrates of different stiffness and across different structural organizations of collagen, including on random collagen fibers, aligned collagen fibers and on isotropically adsorbed collagen. We assess the role of formins and Arp2/3 in controlling traction stresses on aligned collagen fibers. Finally, we determine the role of traction stress in controlling contact guidance across different stiffnesses and in different cell lines. This work describes the biophysical mechanisms that governs cellular mechanotransduction on aligned collagen fiber networks that lead to contact guidance.

## 2. Results

### 2.1. Dual-traction force microscopy can measure deformation of aligned collagen fiber networks and reveals differences in strain fields as a function of depth

We developed a dual-traction force microscopy (d-TFM) approach to assess traction forces at two depths simultaneously. These depths were probed by fluorescent beads embedded within the flexible substrate (TFM1 in **Schematic 1**) and fluorescent beads attached to aligned collagen fibers (TFM2 in **Schematic 1**). Three steps were crucial to carry out d-TFM: (1) embedding of 200 nm fluorescence beads in flexible substrates, (2) transferring of aligned collagen fibers assembled on mica to the top of the flexible substrate and (3) absorbing 40 nm fluorescence beads to aligned collagen fiber networks. We also adsorbed 40 nm beads on aligned collagen fibers assembled on mica and on aligned collagen fibers assembled on mica and transferred to glass ^35^ (**Figure S1**). While the beads were easily adsorbed to the aligned collagen fibers on mica, adsorbing them to collagen fibers transferred to glass and flexible substrates required stringent washing of the gelatin that remained on the fibers after transfer. Improper washing resulted in beads that de-adsorbed once the gelatin was melted away at 37 °C (**Figure S2**). **Schematic 1** compares the d-TFM system with a conventional TFM on several different types of ECM: collagen heterotrimers, which constitute the monomer of the collagen fiber (MM), aligned collagen fibers (AF) and random collagen fibers (RF). Cells can then be plated on these substrates and traction forces on MM, AF and RF can be assessed both in the cell-fiber plane and in the underlying flexible substrate.

Our lab has extensive experience with using mica to create various collagen networks for studying cell migration ^35,36,38^. Collagen monomers in solution adsorb onto the mica surface by means of electrostatic interactions, van der Waals forces and hydrogen bonding ^39^. Electrostatic interactions and a steric ridge influence the orientation of the growing collagen fiber, promoting a parallel alignment, which is critical for generating aligned fiber networks. Intermolecular cross-links form between adjacent collagen molecules, stabilizing the structure and enhancing mechanical strength. AFM imaging reveals collagen alignment when assembled on mica (**Figure 1A**) and when assembled on mica and transferred to glass (**Figure 1B**). The white dotted circle and red solid circle represent the diffraction-limited and actual size of 40 nm fluorescent beads adsorbed to the aligned collagen fibers. High-resolution immunostained aligned collagen fibers show similar aligned structures and can be overlaid with beads embedded in a 2000 Pa PAA gel (BEG) and beads adsorbed on aligned fibers (BAF) transferred to the 2000 pa PAA gel (**Figures 1C&D**). MDA-MB-231 cells readily align on the aligned collagen fibers transferred to 2000 Pa PAA gels (**Figure 1E**), demonstrating that aligned collagen fibers act as a robust contact guidance cue.

**Figure 1:**
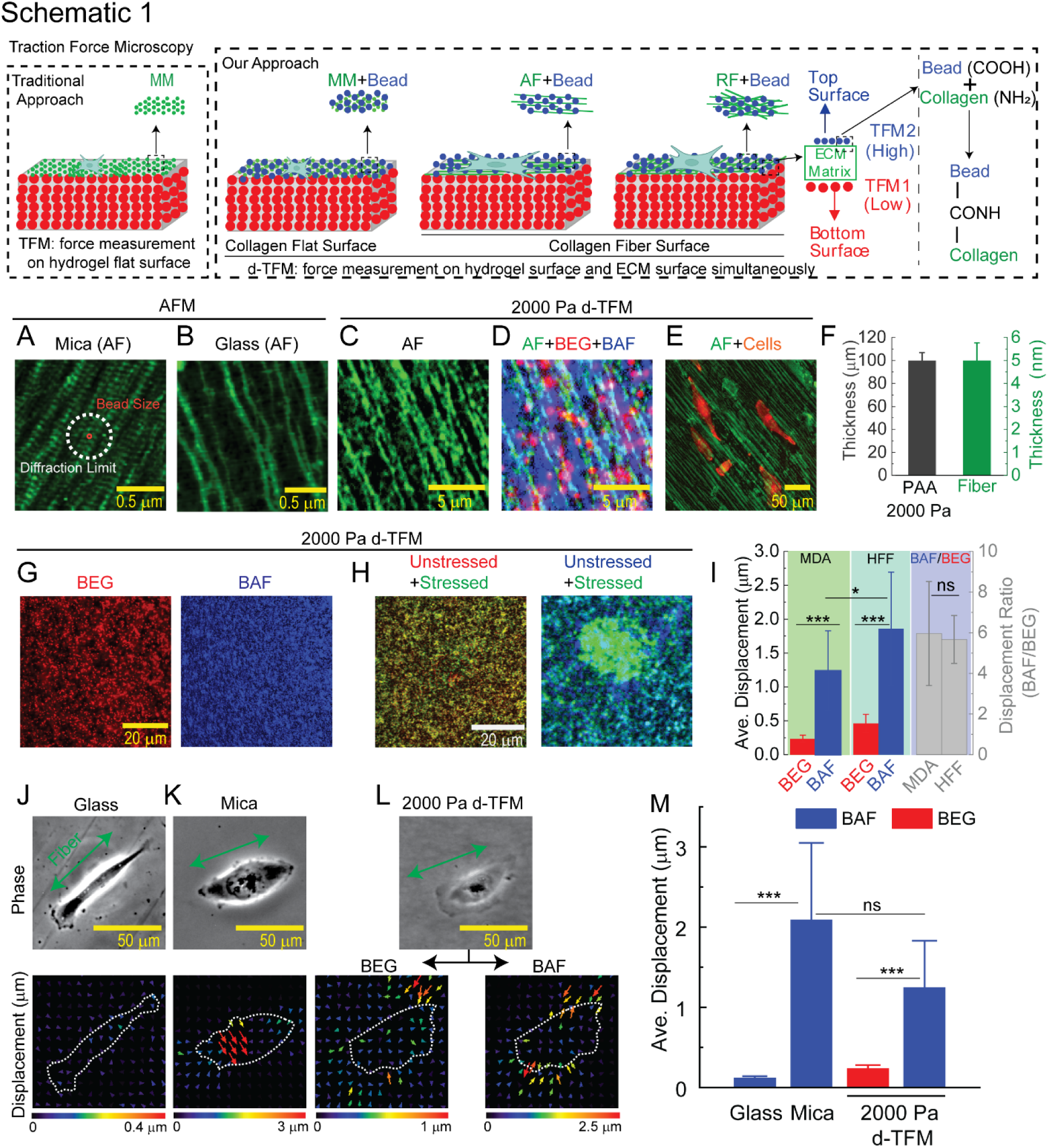
Demonstration of strain field measurement on collagen fiber networks. (**Schematic 1**) Schematic outlining d-TFM on fibers compared to traditional TFM on adsorbed ECM. (**A**) AFM image of collagen fibers assembled on mica where white dotted circle represents diffraction limit of light and solid red circle represents the size of beads attached on fibers used as fiduciary markers. (**B**) AFM image of transferred fibers onto glass from mica. (**C**) High resolution fluorescence image of aligned fibers transferred to a flexible substrate (2000 Pa d-TFM PAA). (**D**) Overlay image of aligned collagen fibers, beads embedded in the PAA gel and beads adsorbed to the fibers on 2000 Pa d-TFM. (**E**) Demonstration of cell alignment along the collagen fibers. (**F**) Thickness of flexible substrate (PAA) and fibers used in our d-TFM system. *N_points, PAA_ = 30 and N_fibers_ = 40*. (**G**) Bead distribution in 2000 Pa PAA and on top of aligned fiber networks (2000 Pa d-TFM). (**H**) Representative images of bead displacement before and after stress exerted by MDA-MB-231 cells at 4 h on 2000 Pa d-TFM. (**I**) Quantitative average displacement of beads underneath cells on BAF and BEG due to stress exerted by MDA-MB-231 (*N_cells_ = 28*) and HFF cells (*N_cells_* = 19) at 2 h and their ratio (BAF/BEG) on 2000 Pa d-TFM. Left *y*-axis (black) for cell displacement on BAF and BEG and right *y*-axis (gray) for displacement ratio. MDA represents as MDA-MB-231. (**J-L**) Represent images of cells on different substrates presenting aligned fibers and the corresponding spatial displacement fields exerted by MDA-MB-231 (*N_cells, glass_ = 21*, *N_cells, mica_ = 19* and *N_cells, 2000 Pa d-TFM_ = 28*). Error bars represent 95% confidence interval unless otherwise stated. *p*-values were evaluated by using a two-tailed unpaired student t-test. * represents *p < 0.05*, *** represents *p < 0.005* and not significant (ns) represents *p* > 0.05. All the experiments were repeated at least three times unless otherwise stated.

Since displacements can be influenced by the substrate thickness, particularly in thin substrates ^40^, we selected a polyacrylamide PAA gel thickness beyond which cells are able to sense the underlying glass substrate (∼100 μm). The average thickness of the polyacrylamide flexible substrate is 100 µm, much larger than the average thickness of collagen fibers, which are 5 nm (**Figure 1F**). We optimized the BAF and BEG bead density to ensure we can track displacements at high resolution, creating bead densities of about 1/μm^2^ (**Figures 1G** and **Figure S3**). MDA-MB-231 cells adhered to collagen fibers transferred to flexible substrates cause substrate deformation, displacing beads underneath (**Figure 1H**). The displacement of collagen fibers attached to 2000 Pa flexible substrates by MDA-MB-231 and human foreskin fibroblast (HFF) cells were analyzed on d-TFM after 2 h of cell attachment. The strong contractile nature of HFFs caused greater displacement than MDA-MB-231s (**Figure 1I**) ^19^. BAF displacement surpassed BEG displacement in both MDA-MB-231s and HFFs. The displacement ratio between BAF and BEG in both MDA-MB-231s and HFFs was identical, suggesting that the strain fields are functions of depth and mechanical coupling of the fibers to the flexible substrate and are not cell-type dependent (**Figure 1I**). This establishes our ability to measure deformations at different depths on aligned collagen fibers transferred to flexible substrates.

Given our ability to measure substrate deformations on top and underneath transferred aligned collagen fibers, we wondered whether the deformation on aligned collagen fibers varied as a function of underlying substrate modulus. The deformations on aligned collagen fibers were measured by quantifying average bead displacement underneath the cell on three different substrates: collagen fibers assembled on mica, assembled on mica and transferred to glass and assembled on mica and transferred to 2000 Pa d-TFM substrates. MDA-MB-231 cells were plated on aligned collagen fibers of the three surfaces and incubated for 15 min prior to recording time-lapse imaging for 2 h (**Figure 1J-L**). Cells deform aligned collagen fibers assembled on mica as we have shown previously ^24^. Additionally, the average displacement of collagen fibers on mica appears to be similar to that observed when collagen fibers are transferred to 2000 Pa d-TFM substrates, suggesting that the observed elastic modulus of collagen fibers assembled on mica is close to 2000 Pa (**Figure 1M**). Few bead deformations were detected on collagen fibers transferred to glass, revealing that beads were strongly attached to aligned collagen fibers and aligned collagen fibers were strongly attached to glass. This establishes our ability to form d-TFM substrates and measure displacements both on top and underneath the cells, revealing that while mica is extremely stiff, collagen fibers assembled on mica form weak interactions with the mica and can be deformed to the same extend as fibers transferred to 2000 Pa d-TFM substrates.

### 2.2 Elastic modulus determines the steady-state level, but not the kinetics of traction stress on aligned collagen fibers

We previously demonstrated that MDA-MB-231 cells spread and align on the three distinct surfaces: collagen fibers assembled on mica, assembled on mica and transferred to glass and assembled on mica and transferred to 2000 Pa d-TFM substrates. We also quantified the deformation. To further characterize traction force behavior on aligned collagen fibers, we calculated traction stress maps for both HFF and MDA-MB-231 cells on different d-TFM substrates as well as explored the kinetics of traction stress development on aligned collagen fibers during spreading. First, we assessed deformations of aligned collagen fibers assembled on mica. HFF and MDA-MB-231 cells were plated on aligned collagen fibers assembled on mica and incubated for 15 min prior to carrying out an 8-h time-lapse imaging experiment. MDA-MB-231 cells remained aligned and migrated on mica for 8 h, whereas HFFs delaminated the collagen fibers from the surface after about 2 h, demonstrating the stronger contractile ability of HFFs (**Figure 2A-B&E** and **Figure S4**). The pronounced contractility of HFFs as the driver of collagen delamination was confirmed by blocking myosin II with blebbistatin. A submaximal concentration of 10 µM blebbistatin effectively diminished the contractile force to a level comparable to that of MDA-MB-231, thereby allowing HFFs to spread and transmit traction force, but inhibiting collagen delamination for up to 8 h (**Figure 2E**). The submaximal concentration of 10 µM blebbistatin resulted in little change in HFF cell area, which was still almost two-fold higher than that of MDA-MB-231 cells (**Figure 2F**). Consequently, myosin II is critical for contractility on aligned collagen fibers and cell area is a poor predictor of traction stress on aligned collagen fiber networks.

**Figure 2:**
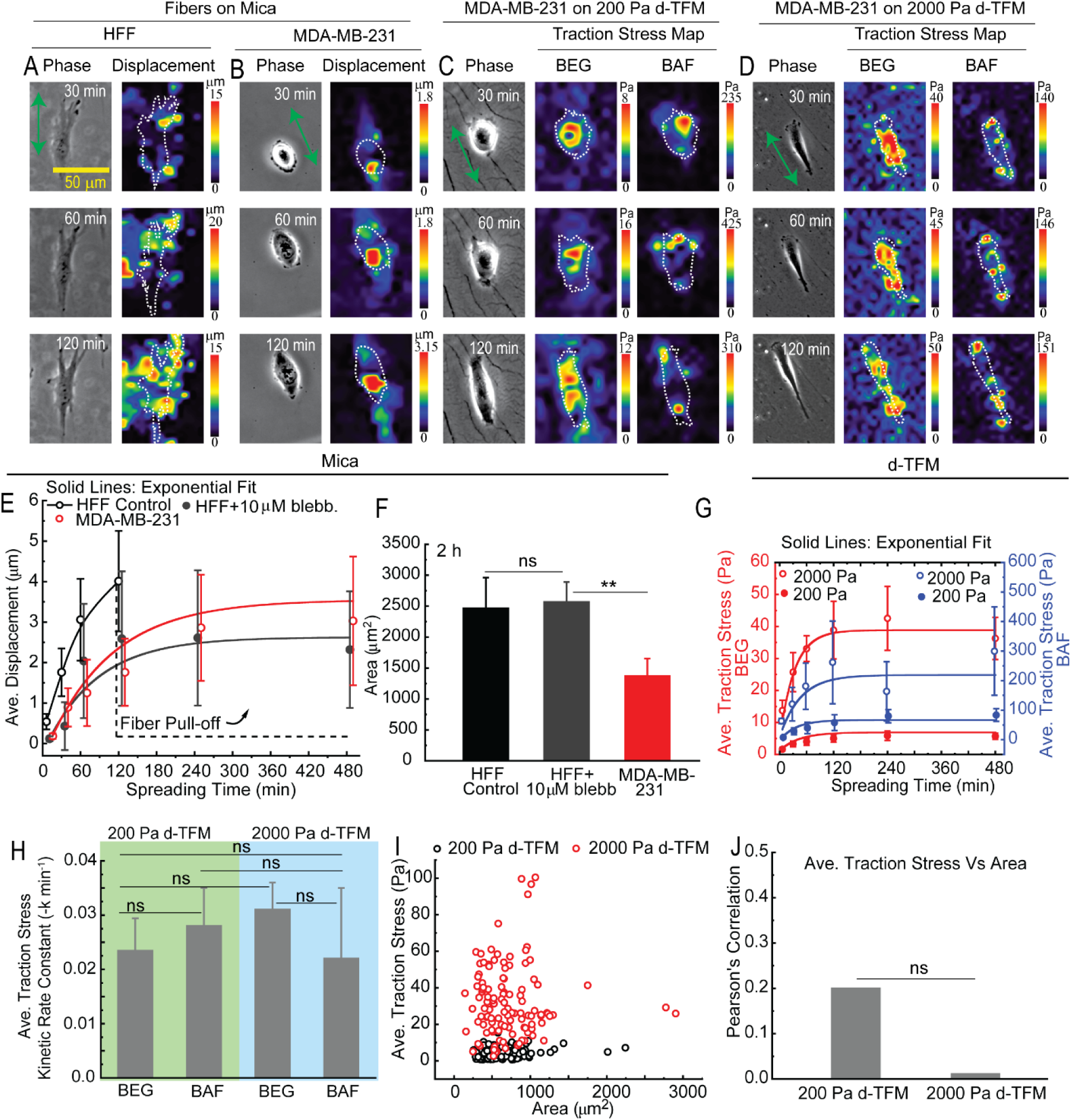
Traction force exerted by HFF and MDA-MB-231 on aligned collagen fibers during spreading. **(A, B)** Displacement of beads exerted by HFF and MDA-MB-231. **(C, D)** Traction stress exerted by MDA-MB-231 on aligned collagen fiber networks attached to 200 Pa d-TFM and 2000 Pa d-TFM PAA gels. **(E)** Displacement kinetics of HFFs, HFFs treated with blebbistatin and MDA-MB-231s. **(F)** Cell area of HFF, HFF treated with blebbistatin and MDA-MB-231 at 2 h. *N_cells, HFF_* = 22, *N_cells, HFF+blebb._* = 19 and *N_cells, MDA-MB-231_* = 19. **(G)** Kinetics of traction stress exerted by MDA-MB-231 on aligned fiber networks using 200 Pa d-TFM (*N_cells_* = 20) and 2000 Pa d-TFM (*N_cells_* = 23). **(H)** Traction stress kinetic constant (*k*, min^-1^) of at each condition determined from fit in G. Error bar represents standard error. **(I)** Correlation between traction stress (from beads embedded in PAA gels, BEG) and area at different spreading times (*N_cells, 200Pa d-TFM_* = 120 and *N_cells, 2000Pa d-TFM_* = 133). **(J)** Pearson’s correlation coefficient between traction stress and area on aligned collagen fiber networks. Error bars represent 95% confidence interval unless otherwise stated unless otherwise stated. *p*-values were evaluated by using a two-tailed unpaired student t-test. ** represents 0.05 < *p <* 0.1, *** represents *p <* 0.005 and not significant (ns) represents *p* > 0.05. All the experiments were replicated at least three times unless otherwise stated.

Next, we conducted time-lapse imaging of MDA-MB-231 on 200 Pa and 2000 Pa d-TFM substrates to quantitatively assess traction stress characteristics. Since the elastic modulus of the underlying substrate is known and we established that the mechanical coupling is constant (**Figure 1I**), traction stress can be calculated on the aligned collagen fibers transferred to d-TFM substrates, unlike the aligned collagen fibers assembled on mica. Phase contrast microscopy images and traction force maps obtained from d-TFM during a duration of 2 h are shown (**Figure 2C&D**). MDA-MB-231 spread slower and align to lesser extents on aligned fibers transferred to 200 Pa substrates than on aligned fibers transferred to 2000 Pa substrates. As cells spread, they transmit higher average traction stress (**Figure 2G**), agreeing with previous work indicating that contractility increases during spreading leading to higher traction forces ^41^. As described above for deformations, the exerted traction stress transmitted on the BEG (TFM1) surface were substantially lower than the force exerted on the BAF surface (TFM2) over all times. These results show that higher stresses are exerted on aligned collagen fiber surfaces as compared to the underlying substrate, regardless of elastic modulus (200 Pa and 2000 Pa). The steady-state level of traction stress on aligned collagen fibers increased with increasing elastic modulus as is commonly observed in non-fiber ECM systems. Unlike non-fiber ECM systems, the spread area is not a good predictor of average traction stress exerted whether on 200 Pa or 2000 Pa substrates (**Figure 2I&J**). Finally, the average traction stress kinetics reported using BEG displacements and BAF displacements were fitted well with a saturating exponential function and the kinetic rate constant was found to be roughly similar, varying much less than the differences in magnitude (**Figure 2H**). This reveals that while the magnitude of traction stress is stiffness-dependent on aligned collagen fibers, the kinetics are not.

To assess the cell-to-cell variability in the application of d-TFM in reporting traction stress on aligned collagen fibers, we examined multiple MDA-MB-231 cells on 2000 Pa d-TFM substrates (**Figure 3A**). Greater variability in traction stress maps were reported by BAF as compared to BEG (**Figure 3A&C**). A possible explanation of this higher variability is that cells exert large gradients in strain on the fibers, resulting in localized regions with a small population of large bead displacements. The underlying substrate both dampens the magnitude and smooths this spatial heterogeneity through stress dissipated by the flexible substrate as a function of depth, resulting in less variation. Additionally, traction stress exerted by MDA-MB-231 cells was measured on d-TFM substrates of 200 Pa, 2000 Pa and 20,000 Pa (**Figure 3B&D**). Higher traction stress was calculated using BAF displacements as compared to BEG displacements. Additionally, both average traction forces calculated using BAF and BEG displacements increased with increasing elastic modulus. Overall, cells exert heterogeneous traction forces on aligned collagen fibers transferred to d-TFM substrates, but the mean cell response shows increasing average traction force increases as a function of elastic modulus as described for non-fiber systems.

**Figure 3:**
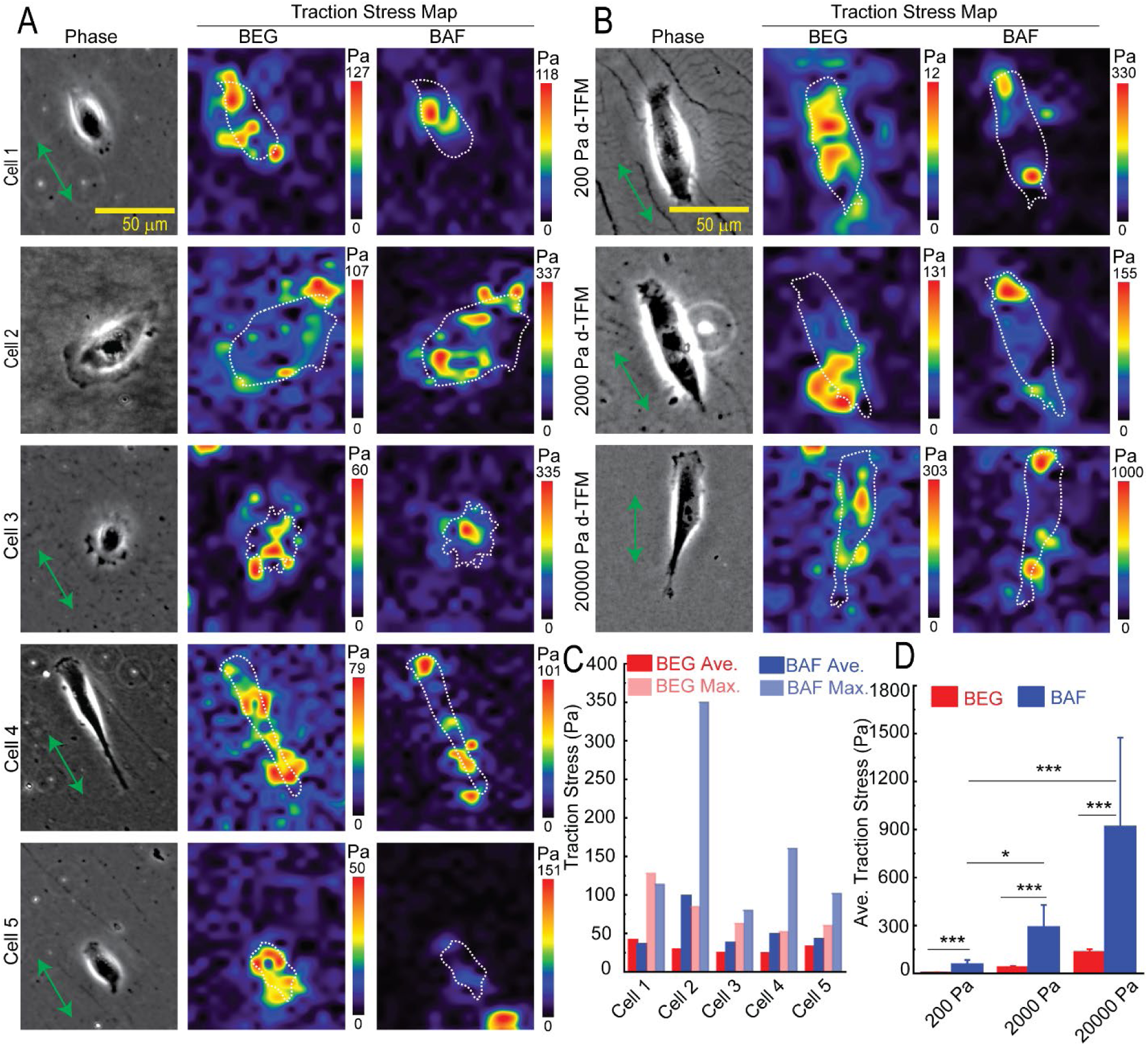
Heterogeneity of traction forces exerted by MDA-MB-231 on aligned collagen fibers after spreading. **(A)** Representative images of traction stress distribution in different cells on aligned collagen fiber networks on 2000 Pa d-TFM. **(B)** Traction stress on aligned collagen fiber networks on 200 Pa d-TFM, 2000 Pa d-TFM and 20000 Pa d-TFM. **(C)** Traction stress variation in each cell corresponding to **(A) (***N_cells_* = 5). **(D)** Stiffness-dependent variation in traction stress corresponding to BEG and BAF on 200 Pa d-TFM (*N_cells_* = 20), 2000 Pa d-TFM (*N_cells_* = 23) and 20000 Pa d-TFM (*N_cells_* = 23) **(B)**. Error bars represent 95% confidence interval unless otherwise stated. *p*-values were evaluated by using a two-tailed unpaired student t-test. *represents *p < 0.05* and *** represents *p < 0.005*. All the experiments were replicated at least three times unless otherwise stated.

### 2.3 The structure of the collagen network determines the magnitude and the kinetics of traction stress

Given that we characterized displacements on aligned collagen fibers assembled on mica and traction stresses on aligned collagen fibers transferred to d-TFM substrates, we were interested in determining how the traction stress magnitudes and kinetics for homogeneously adsorbed collagen monomers compared to aligned and randomly oriented collagen fibers. We have shown that higher collagen adsorption concentrations result in random fiber deposition on mica and non-directional migration ^36^. Thus, we were able to probe three distinct ECM environments on 2000 Pa substrates: isotropic collagen monomers (MM), anisotropic aligned collagen fibers (AF) and isotropic random collagen fibers (RF) by adsorbing collagen monomers onto the flexible substrate as others have done and using our approach to transfer either aligned or random collagen fibers to 2000 Pa d-TFM substrates. The three types of adsorption caused different collagen staining patterns across the three distinct substrates (**Figures 4A-C**). Collagen staining intensity varied somewhat between the three different substrates, where monomers (MM) exhibited a lower intensity than either of the fiber networks (AF and RF) (**Figure 4D**). The higher intensity level of fiber networks is likely due to the structurally organized and well packed interconnected pattern of fibers. HFF cells were incubated on MM, AF, and RF on d-TFM at 2000 Pa for 15 min to allow them to adhere first, and then the images were taken for 8 h. The resulting phase contrast image and average traction stress map on BEG (beads embedded in PAA gel) and BMM (beads attached on monomers), BAF (beads attached on aligned fibers), and BRF (beads attached on random fibers) surfaces on d-TFM at 2000 Pa of each substrate are shown (**Figures 4E-G**). As expected, we found that traction force exerted on BMM, BAF, and BRF clearly surpassed the force exerted on BEG, i.e., TFM2 (BMM, BAF, BRF) > TFM1 (BEG) under each collagen structural condition (**Figure 4H-J**, maximum traction force is included in **Figure S5A-C**), suggesting that a traction force reported in previous published work likely underestimates traction force due to a dissipation of displacement as a function of depth. Indeed, the average traction stress ratio between TFM1 and TFM2 is nearly 8 for each condition: MM, AF, and RF (**Figure 4N**). The traction stress correlation between TFM1 and TFM2 reveals a moderate correlation regardless of collagen structure (**Figure S5G-I**). Traction forces are higher on collagen fiber networks (AF or RF) as compared to the isotropic collagen surfaces (MM) starting vary early during spreading.

**Figure 4:**
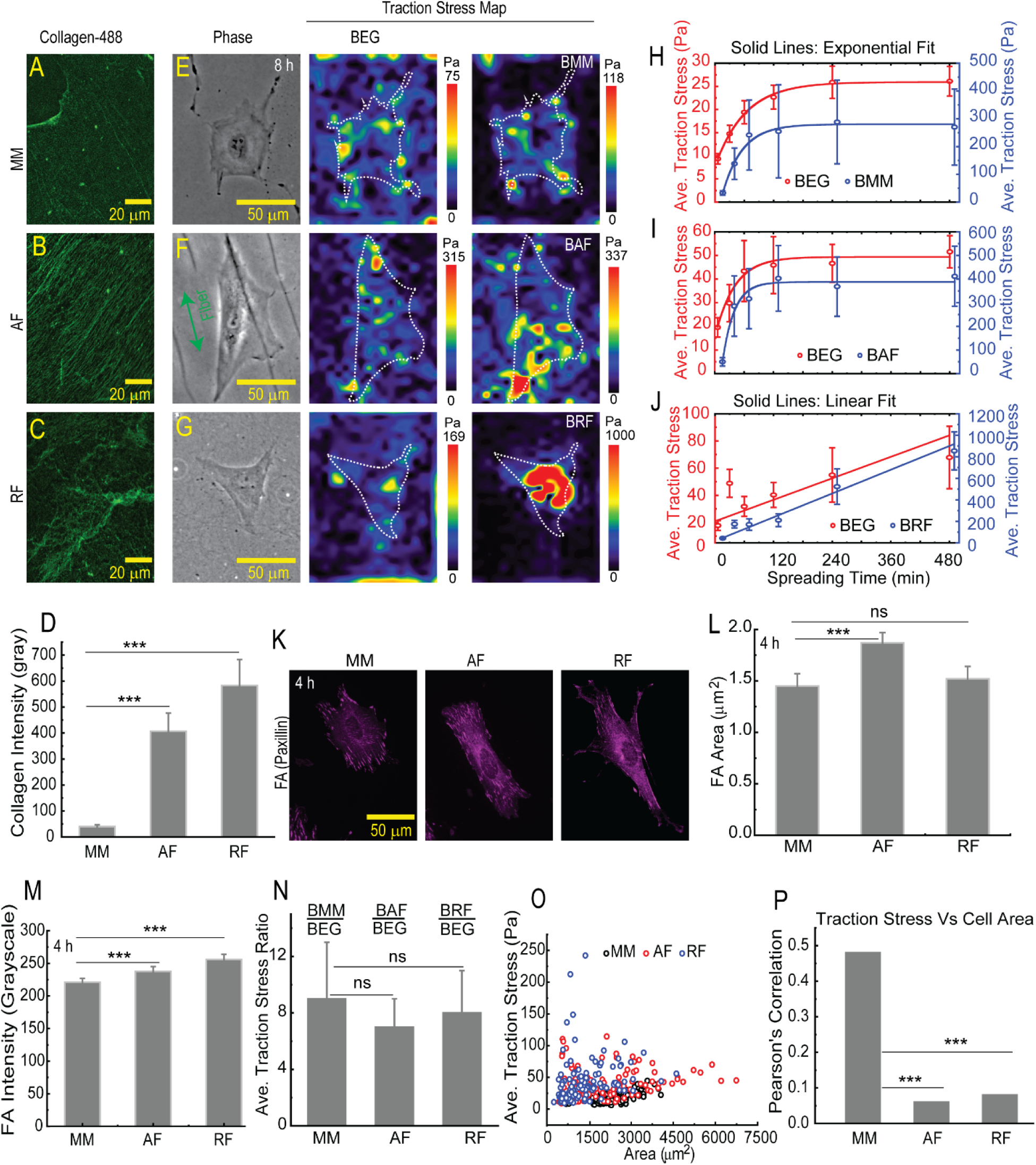
Traction force exerted by HFFs on different collagen networks. **(A-C)** Immunofluorescence images of collagen monomers (MM), transferred aligned collagen fibers (AF) and random collagen fibers (RF) on 2000 Pa d-TFM substrates. **(D)** Quantified immunofluorescence intensity corresponding MM, AF and RF respectively. *(N_points, MM_* = 29*, N_points, AF_* = 27 *and N_points, RF_* = 25. (**E-G**) Phase and traction stress map exerted by HFF on distinct structural cues MM, AF and RF on 2000 Pa d-TFM. **(H-J)** Traction stress kinetics corresponding to **(E-G)** respectively. *N_cells, MM_* = 20, *N_cells, AF_* = 19 and *N_cells, RF_*= 20. (**K**) Paxillin stained fluorescence images on distinct structural cues MM, AF and RF on 2000 Pa d-TFM**. (L**) FA size on MM, AF and RF on 2000 Pa d-TFM at 4 h. *N_points,_ MM, AF, RF* ≥ 200. (M) Fluorescence intensity of FAs (paxillin) stained HFFs on MM, AF and RF on 2000 Pa d-TFM at 4 h, corresponding to K and L. (**N**) Average ratiometric traction stress exerted by BEG and BMM, BAF, BRF on 2000 Pa d-TFM respectively within 8 h. **(O)** Distribution of traction stress with respect to cell area on MM, AF and RF on2000 Pa d-TFM at different time points. *N_cells, MM_ = 120, N_cells, AF_ = 114 and N_cells, RF_= 120*. Here, we correlated the traction stress exerted on BEG surface. **(P)** Pearson’s correlation coefficient between traction stress and cell area corresponding to MM, AF and RF on 2000 Pa d-TFM. Error bars represent 95% confidence interval unless otherwise stated. *P*-values were evaluated by using a two-tailed unpaired student t-test. *** represents *p < 0.005* and ns represents non-significant. All the experiments were replicated at least three times unless otherwise stated.

This occurs even though the spread areas are approximately similar (**Figure 4O** and **Figure S5D-F**). Surprisingly, we observed very different traction stress kinetics when comparing AF and RF surfaces (**Figure 4H-J** and **Figure S5A-C**). While AF surfaces result in an exponential growth of the traction stress similar to what is seen on MM surfaces, coming to a steady-state around 2 h, RF surfaces show roughly linear increases in traction stress over 8 h. To determine if these changes in cell traction produce changes in adhesion, FAs were stained (**Figure 4K**) and focal adhesion size and intensity was quantified at 4 h (**Figure 4L&M**). The FA area on AF is marginally larger than on MM and RF surfaces, indicating a lack of direct correlation between the FA area and the magnitude of the traction force (**Figure 4L**).

However, RF substrate exhibited higher FA intensity, indicating stronger adhesive strength and, consequently, greater traction force transmission compared to AF and MM (**Figure 4M**). We also examined the traction force correlation with the cellular area (**Figure 4O&P** and **Figure S6D-F**). Traction force is highly correlated with cell area for isotropic adsorbed collagen (**Figure 4O&P**) ^42,43^. However, traction force is completely uncorrelated with cell area for either aligned or random fibers (**Figure 4O&P**). Consequently, there are distinct traction force signatures that collagen fibers elicit in comparison to isotropically adsorbed collagen.

### 2.4. Formin and Arp2/3 diminish maximum traction stress, while not affecting the average traction stress in mesenchymal cells on aligned collagen fibers

We demonstrated that aligned collagen fibers result in unique traction stress signatures as compared to collagen monomers. It is known that F-actin is a required for traction force generation, and inhibition of F-actin polymerization significantly reduces traction force on isotropic ECM ^44–46^. However, F-actin can be assembled either into linear networks through formins or branched networks through Arp2/3. Formins produce linear F-actin filaments termed stress fibers that can be aligned in the direction of collagen fiber alignment and generate contractile force, facilitating contact guidance. Arp2/3 produces branched F-actin filaments that might allow for cell turning away from the fiber direction through the generation of protrusive force, diminishing contact guidance ^47^. While there is some evidence that formins positively regulate contact guidance, Arp2/3 has been shown to have positive ^48^, negative or no affect on contact guidance ^27^. We wondered how formin and Arp2/3 contribute to the average and maximum traction force on aligned collagen fibers. We treated a mesenchymal cell line (HFFs) that have relatively high contact guidance fidelity with inhibitors for formins (SMIFH2) and Arp2/3 (CK-666). In addition, we explored knocked down expression of a particular formin, Formin homology 2 domain-containing 3 (FHOD3) protein using siRNA. FHOD3 was selected as it is known to regulate contact guidance ^48,49^. We hypothesized that formins, regulators of stress fibers, rather than Arp2/3, a regulator of protrusion and not contraction, would more strongly control traction stress generation on aligned collagen fibers.

HFFs treated with SMIFH2 or CK-666 were plated on aligned collagen fibers on d-TFM at 2000 Pa. Cells were allowed to spread for 15 min followed by 4 h of imaging (**Figure 5A**). Myosin II inhibition occurs at high concentrations of SMIFH2 (> 50 μM), consequently, we used 20 μM SMIFH2, a lower concentration that should not affect myosin II activity ^50^. To supplement pharmacological inhibition and to explore the role of a specific formin, we used siRNA to knock down FHOD3 transcript levels.

**Figure 5:**
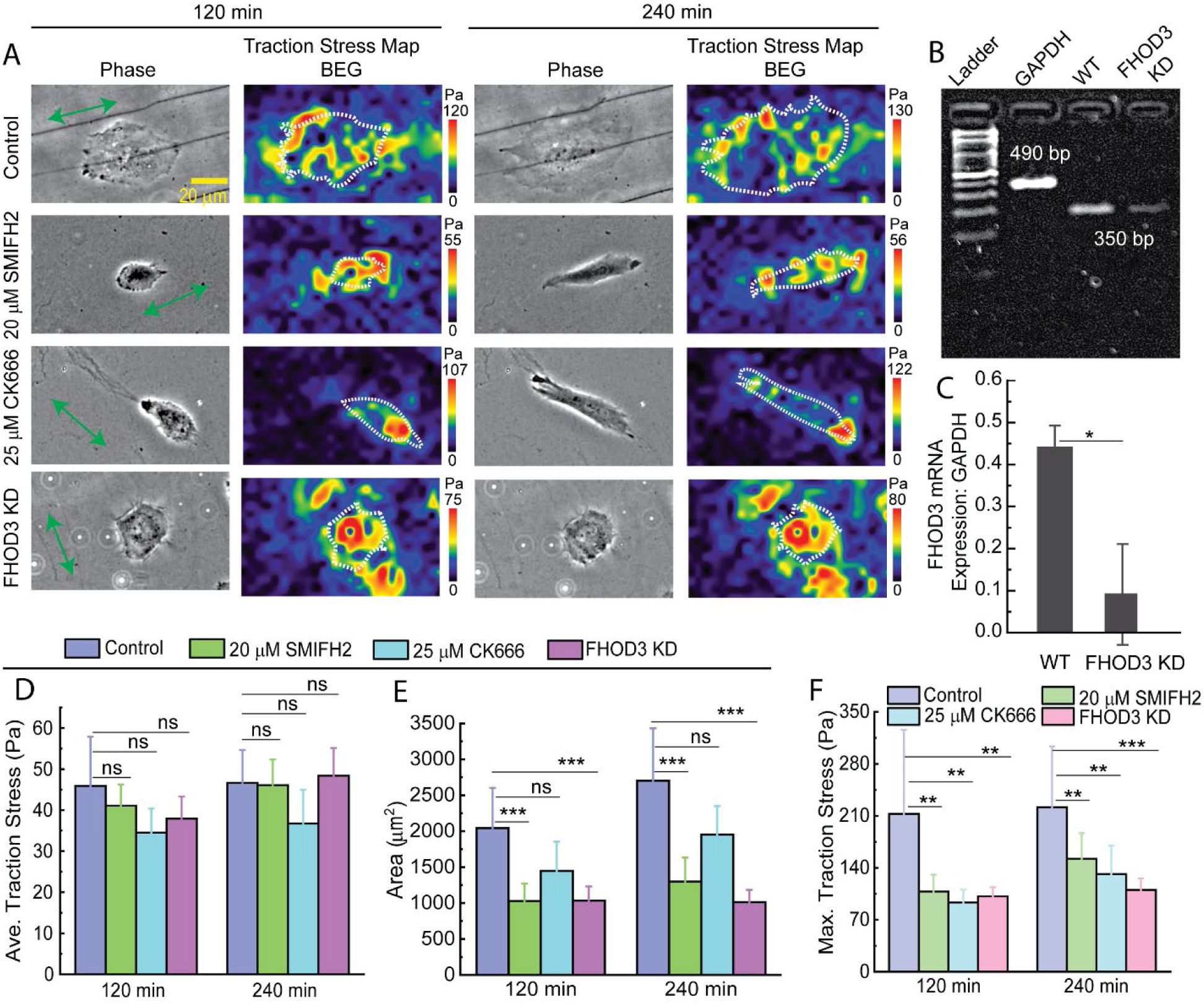
The role of formins and Arp2/3 on traction force exerted by HFF on aligned collagen fibers networks. (**A**) Phase contrast microscopy and traction stress map on 2000 Pa d-TFM at 2 h and 4 h. **(B)** Formin expression measured by RT-PCR. (**C**) FHOD3 mRNA expression level of WT and after FHOD3 knock-down (FHOD3 KD). *N_samples_* = 2. (**D-F**) Average traction stress, average cell area and maximum traction stress under different condition corresponding to **(A)**. *N_cells, control_ = 19, N_cells, SMIFH2_ = 25, N_cells, CK-666_= 19 and N_cells, FHOD3 KD_ = 20*. Error bars represent 95% confidence interval unless otherwise stated *P*-values were evaluated by using a two-tailed unpaired student t-test. * represents *p < 0.05,* ** represents 0.05 < *p < 0.1, \*\*\** represents *p < 0.005* and ns represents non-significant. All the experiments were replicated at least three times unless otherwise stated.

Treatment with siRNA resulted in an 80% knock-down of the transcript levels in HFFs (**Figure 5B&C**). Inhibiting Arp2/3 using CK-666, inhibiting formins using SMIFH2 or knocking down FHOD3 resulted in no change in average traction stress (**Figure 5D**). However, inhibiting formins with SMIFH2 as well as knocking down FHOD3 significantly diminished cell area (**Figure 5E**). No change in cell area was seen after Arp2/3 inhibition. The response to both formin inhibitor and knockdown of FHOD3 were similar, suggesting that FHOD3 is the primary formin that drives changes in area. Additionally, all perturbations resulted in a diminished maximum traction stress (**Figure 5F**). Consequently, while formins and Arp2/3 do not regulate average traction stress, both affect the maximum traction stress. Formins and in particular FHOD3 facilitates spreading. This indicates that both formins and Arp2/3 are required for traction force generation on aligned collagen fibers, but they have distinct roles in controlling the spread area of mesenchymal cells.

### 2.5. Formin and Arp2/3 are both necessary for high force transmission during epithelial cell turning, but formins regulate the traction force magnitude, whereas Arp2/3 controls the traction force kinetics on aligned collagen fibers

We showed that formins and Arp2/3 appear to both be important in regulating traction stress in mesenchymal cells. However, these cells robustly align in the direction of collagen fiber alignment^36,37^. Mesenchymal cells migrate following the fiber direction with high fidelity. Consequently, we were interested in exploring the role of formins and Arp2/3 in regulating traction stress in cells that do not follow aligned collagen fibers with high fidelity. Epithelial cells are less contractile and make more frequent turns on aligned fibers. Epithelial cells appear to exhibit two migration strategies, one that is perpendicular and one that is parallel to collagen fiber alignment (paper in revision). The traction stress behavior during cell turning is not known. Furthermore, how formins and Arp2/3 regulate the turning behavior of cells is also unknown. We choose a human keratinocyte cell line (HaCaT) that exhibits this turning behavior as a representative epithelial cell line for our study. We examined the force transmission characteristics during cell turning from parallel to perpendicular and vice versa with respect to aligned collagen fibers.

We plated HaCaT cells onto aligned collagen fibers on d-TFM substrates at 2000 Pa, incubated them for 15 min to allow them to adhere, and then recorded a time-lapse video for 8 h. We measured the traction force of the cells, classifying them into three types of alignment with respect to the aligned collagen fibers (**Figure 6A**). Cell direction was defined as parallel to the aligned collagen fibers if 0° ≤ *θ* < 30°, transitional to the aligned collagen fibers if 30° ≤ *θ* < 60° and perpendicular to the aligned collagen fibers if 60° ≤ *θ* ≤ 90°, where *θ* is the angle between the collagen fiber direction and the cell alignment direction (**Figure 6B**). Some HaCaT cells transition between parallel and perpendicular alignment, resulting in changes in traction stress (**Figure 6C**). We observe that when most cells turn from parallel or perpendicular to a transition angle, the average traction stress increases (**Figure 6D**). This is shown with the greyed areas. Next, we measured the traction stress as a function of cell alignment with respect to aligned collagen fibers (**Figure 6E** and **Figure S7B**). The results showed when cells are moving parallel to collagen fiber alignment, the average traction stress is marginally higher than the average traction stress when cells are moving perpendicular to collagen fiber alignment. However, greatest traction stress was achieved at transition angles between parallel and perpendicular movement, around 30° to 45° (**Figure 6E** and **Figure S7B**). Cells treated with SMIFH2 or CK-666 for 8 h reduced this transitional traction stress barrier, highlighting the crucial role of formins and Arp2/3 in generating a higher traction force during cell turning (**Figure 6E**). In addition, we examined the cell population distribution in terms of alignment direction (**Figure 6F**). The highest percentage of cells were aligned parallel to the aligned collagen fibers (∼30%). However, blocking Arp2/3 with CK-666 enhances the fraction of cells that are aligned (15°-25°) with the collagen fibers, indicating that Arp2/3 activity confuses cell alignment by enhancing cell turning on aligned collagen fibers. There was also a significant portion of control cells that were aligned at angles between 30°-45° with respect to the aligned collagen fibers, indicating a potential barrier from moving between a parallel and perpendicular mode of migration. Blocking formins or Arp2/3 diminished the number of cells stuck in this high traction stress, transitional angle. Neither SMIFH2 nor CK-666 affected perpendicular migration (**Figure 6F**). Taken together Arp2/3 acts to diminish parallel migration and both formins and Arp2/3 regulate the high traction stress state associated with epithelial cell turning.

**Figure 6:**
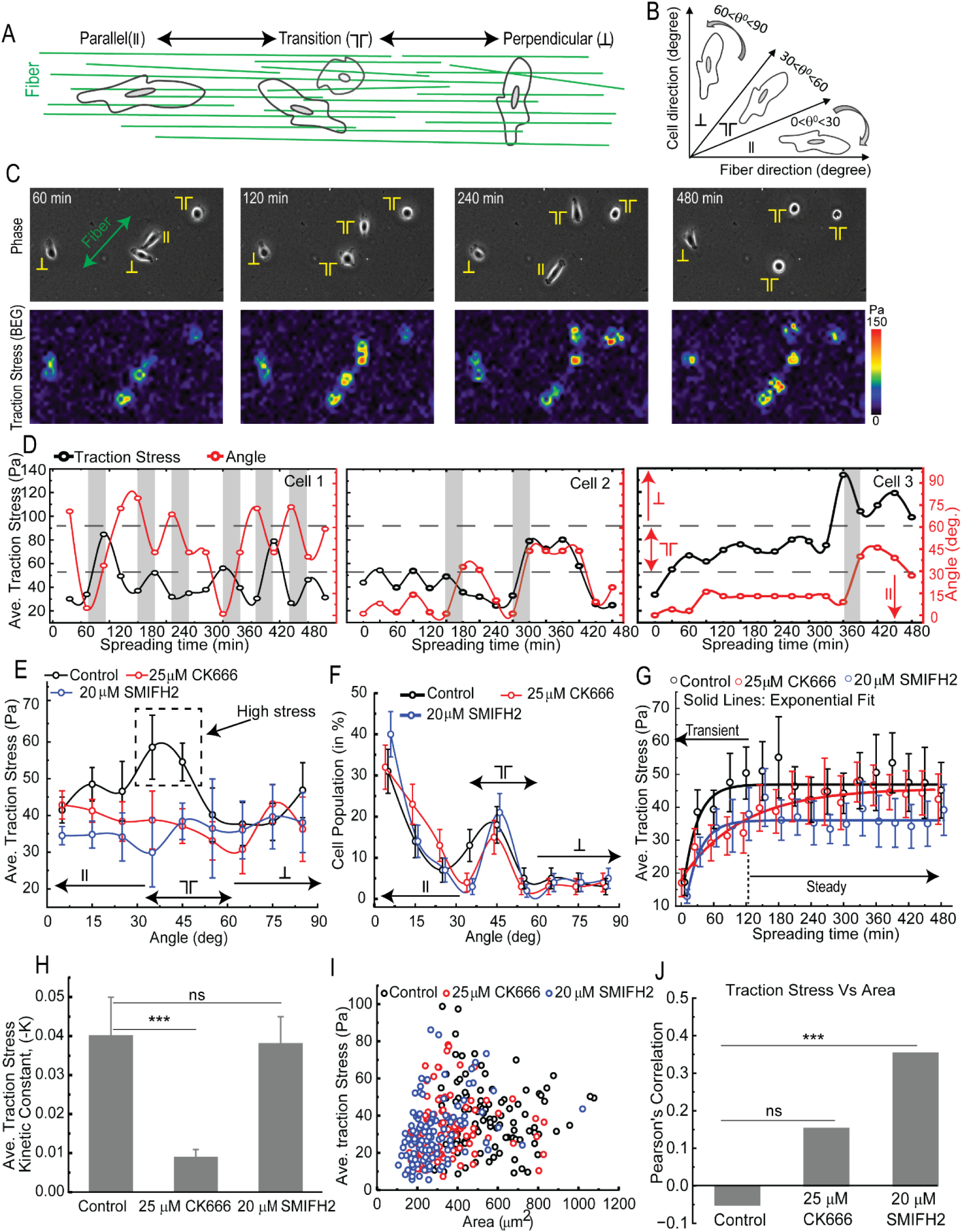
The role of formins and Arp2/3 on traction force exerted by HaCaTs on aligned collagen fibers networks. **(A and B)** Schematic represents three types of cell behaviors in response to aligned fiber networks, i.e., parallel, transition and perpendicular. **(C)** Phase contrast microscopy and traction stress map generated by parallel, transition and perpendicular cells in response to aligned fiber networks. **(D)** Representative cells demonstrating the dynamics of traction stress with respect to spreading time and cell angle on aligned fibers on 2000 Pa d-TFM. (**E**) Traction stress distribution at distinct direction (parallel, transition and perpendicular) under different conditions. *N_cells, control_* = 18, *N_points_* = 288, *N_cells, CK-666_* = 18, *N_points_* = 288 *and N_cells, SMIFH2_* = 19, *N_points_* = 304. (**F**) Cell population density in parallel, transitional and perpendicular orientation across 8 h at the interval of 30 min. in response to aligned collagen fiber networks. **(G)** Kinetics of traction stress (BEG) under different condition in aligned fiber networks on 2000 Pa d-TFM. *N_cells, control_* = 18*, N_cells, CK-666_* = 18 and *N_cells, SMIFH2_* = 19. (**H**) Traction stress kinetic constant (*k, min^-^*^1^) of each condition extracted from fitting kinetics evolution corresponding to E. Error bars represent standard error. **(I)** Distribution of traction stress with respect to cell area at different time. *N_cells, control_ =108, N_cells, CK-666_ =108 and N_cells, SMIFH2_ =114.* **(J)** Represents the Pearson’s correlation coefficient between traction stress and area under different conditions corresponding to G within 8 h. Error bars represent 95% confidence interval unless otherwise stated. *P*-values were evaluated by using a two-tailed unpaired student t-test. *** represents *p < 0.005* and ns represents non-significant. All the experiments were replicated at least three times unless otherwise stated.

Given that both formins and Arp2/3 regulate the high traction stress barrier that occurs at transitional angles, we were interested in whether formins and Arp2/3 regulate the kinetics of traction force generation. The average traction stresses exerted by individual HaCaT cells showed an exponential increase over time, reaching a steady-state level of 45 Pa in about 2 h (**Figure 6G** and **Figure S7A**).

These levels were slightly lower than those observed for HFFs (**Figure 4I** and **Figure 5B**). Formin inhibition resulted in a decrease in the magnitude of traction stress in HaCaT cells, unlike in HFF cells, which showed no decrease in average traction stress. Arp 2/3 resulted in no decrease in the magnitude of traction force, similar to what was found for HFF cells (**Figure 4D**). Rather, inhibiting Arp2/3 in HaCaT cells diminished the rate constant, producing slower traction force kinetics (**Figures 6G&H**). In addition to traction stress kinetics, we explored how these F-actin regulatory proteins affect the correlation between traction stress and spread area (**Figure 6I**). Previously, we established that area is a poor predictor for traction stress on aligned fibers in HFF cells (**Figure 2I&J** and **Figure 4O&P**). There is also no correlation between average traction stress and area in HaCaT cells as exhibited by a Pearson’s correlation coefficient of nearly zero (**Figure 6J**). However, cells treated with SMIFH2 showed an increase in the correlation between traction stress and area, rising to levels that were observed when cells were plated on isotropic collagen substrates rather than collagen fibers. This suggests that blocking formins eliminates the anisotropic force transmission behavior seen on collagen fibers. Taken together, formins and Arp2/3 both regulate the traction stress state during turning. However, Arp2/3 leads to quick changes in average traction stress, whereas formins enhance the magnitude of the average traction stress and is responsible for the uncoupling between average traction stress and area.

### 2.6. Traction force is positively correlated with speed and directionality on aligned collagen fibers

Given the differences in traction stress between random fibers, aligned fibers and uniformly deposited collagen monomers, we were interested in whether traction force determines how well cells engage in contact guidance. Several studies have demonstrated a robust contact guidance on 2D substrates, such as parallel grooves, ridges, or adhesive stripes ^51–53^. Contact guidance along microgrooves requires both cell-substrate adhesions and cellular tension ^54^. However, aligned fiber networks hold greater physiological and pathophysiological significance due to their ability to replicate contact guidance diversity amongst cells in 3D ^55^. Several studies have demonstrated that various cells exhibit distinct contact guidance responses in or on aligned fibers ^19,25,35,56–58^ and high myosin contractility and adhesion strength, required characteristics for high traction stress, are both correlative of strong contact guidance ^24^. However, a direct relationship between contact guidance and traction stress on aligned fibers has yet to be investigated. We hypothesize that increases in traction stress will result in increased contact guidance, but only in the presence of contact guidance cues.

In order to investigate this hypothesis, HFF cells were plated onto three different substrates, MM, AF, and RF, at 2000 Pa (**Figure 7A-C**) for different times and were fixed and stained with phalloidin, a stain for F-actin (**Figures 7D-F** and **Figure S8**). We quantified cell alignment using a directionality index. The directionality index was calculated by finding the average cell alignment direction within one image and using this angle in the directionality index calculation as outlined in the materials and methods.

**Figure 7:**
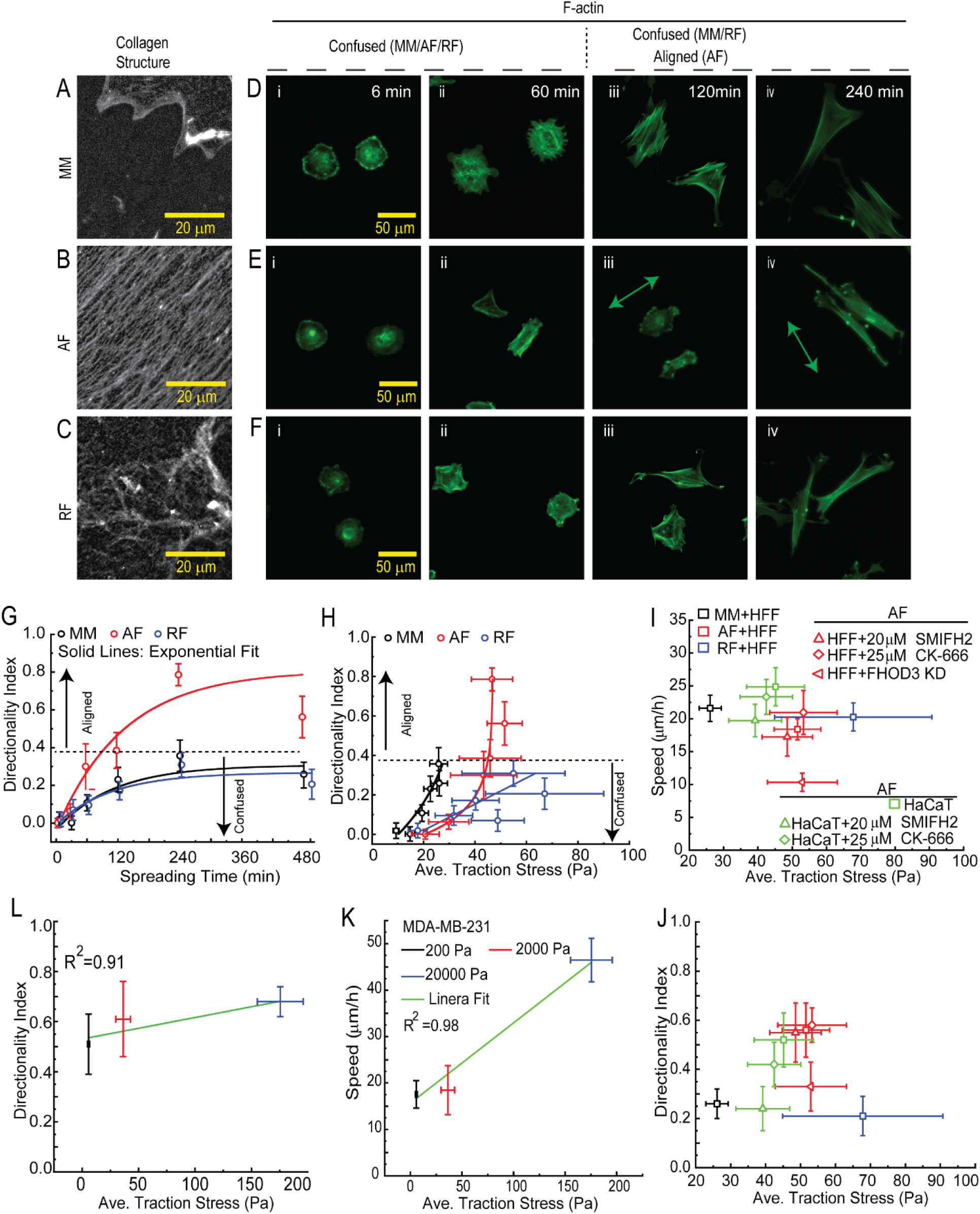
Correlation between traction force, directionality index and speed of cell migration exhibited by different cells on 2000 Pa d-TFM. (**A-C**) Immunofluorescence images of collagen monomers (MM), transferred aligned fibers (AF) and random fibers (RF) on 2000 Pa d-TFM respectively. (**D-F**) Fluorescence images of phalloidin-stained F-actin under different incubation times on distinct structural cues (MM, AF, RF) at 2000 Pa d-TFM respectively. (**G**) Kinetics of directionality index on collagen organizations corresponding to (D-F). *N_cells at each time_* ≥ 143. (**H**) Correlation between traction stress (TS) and directionality index on distinct collagen organizations corresponding to (D-F). MM: *N_cells, TS_* = 20; AF: *N_cells, TS_* = 19 *and* RF: *N_cells, TS_* = 19. (**I**) Correlation between traction stress and speed of cell migration on 2000 Pa d-TFM. HFF (MM: *N_cells, speed_* = 26*, N_cells, TS_* = 20; AF: *N_cells, speed_* = 21*, N_cells, TS_* = 19 and RF: *N_cells, speed_* = 27*, N_cells, TS_* = 19. HFF-AF (Control: *N_cells, speed_* = 26, *N_cells, TS_* = 18; SMIFH2: *N_cells, speed_* = 49, *N_cells, TS_* = 19; CK-666: *N_cells, speed_* = 49, *N_cells, TS_* = 18; FHOD3 KD: *N_cells, speed_* = 29, *N_cells, TS_* = 20*, N_samples_*= 2), HaCaT (Control: *N_cells, speed_* = 20, *N_cells, TS_* = 18; SMIFH2: *N_cells, speed_* = 49, *N_cells, TS_* = 19; CK-666: *N_cells, speed_* = 49, *N_cells, TS_* = 18. (**J**) Correlation between traction stress and directionality index on 2000 Pa d-TFM. HFF (MM: *N_cells, DI_* = 54*, N_cells, TS_* = 20; AF: *N_cells, DI_* = 41*, N_cells, TS_* = 19; HaCaT (Control: *N_cells, DI_*= 47, *N_cells, TS_* = 18; SMIFH2: *N_cells, DI_* = 58, *N_cells, TS_* = 19; CK-666: *N_cells, DI_* = 93, *N_cells, TS_ =* 18 and FHOD3 KD: *N_cells, DI_* = 62, *N_cells, TS_* = 20*, N_samples_*= 2). (**K**) Correlation between traction stress and speed of cell migration on 200 Pa d-TFM, 2000 Pa d-TFM and 20000 Pa d-TFM and (**L**) Correlation between traction stress and directionality index on 200 Pa d-TFM (*N_cells, DI_* = 40, *N_cells, TS_* = 20), 2000 Pa d-TFM (*N_cells, DI_* = 27, *N_cells, TS_* = 23) and 20000 Pa d-TFM (*N_cells, DI_* = 59, *N_cells, TS_* = 62). Error bars represent 95% confidence interval. All the experiments were replicated at least three times unless otherwise stated.

**Figure 8:**
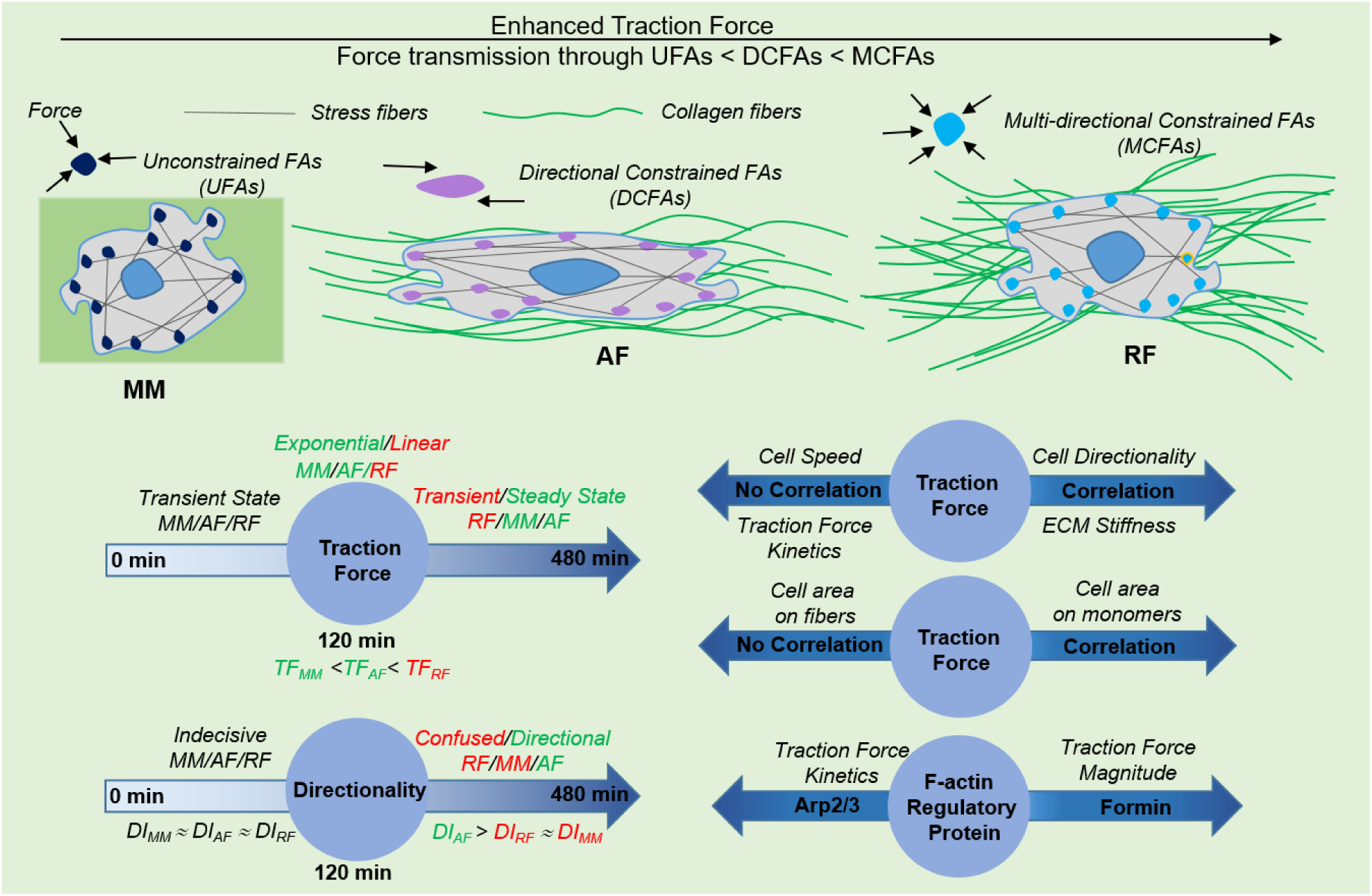
Establishing the general correlation between traction forces, collagen networks and cell behaviors.

Directionality index values of 1 indicate high alignment with collagen fibers and high-fidelity contact guidance, and values of 0 indicate low alignment with collagen fibers and low-fidelity contact guidance. Cells were considered aligned if their directionality index was above 0.4 and unaligned if below 0.4. As expected, the directionality index of cells on MM, AF and RF was almost the same for early adhesion until 30 min (**Figure 7G**). However, distinct changes in directionality index were observed after 1 h. On AF substrates, the directionality index increases robustly, whereas on MM an RF substrates, there is a slow increase to 0.3. Within 2 h, cells on AF substrates make a directional decision whether to be aligned in the direction of collagen fibers or not (**Figure 7D-G)**. On MM and RF substrates, cells are indecisive and orient in different directions over longer timescales (**Figures 7D-G**). The directionality index after 6 h was about double on AF substrates as it was on MM or RF substrates.

Next, we examined the correlation between traction forces and the directionality index on these different substrates. Interestingly, we observed an exponential increase in directionality index as a function of traction stress on AF substrates. MM substrates resulted in only small increases in traction force. RF substrates were more variable and seemed to show more of a linear increase in directionality as a function of traction stress. These results are consistent with the traction force kinetics observed previously (**Figure 4H-J**). It appears that cells require a minimum level of average traction stress (40 Pa) to initiate directional alignment, but once some directional alignment is achieved (*DI* ∼ 0.4) increased traction stress is not needed. Interestingly, RF substrates seem to allow the cell to generate high traction stress, but the lack of collagen fiber alignment does not produce aligned cells. In contrast, MM substrates seem to lack both the high traction stresses associated with alignment as well as cell alignment. Only AF substrates support robust cell alignment by stimulating enough traction stress as well as presenting the aligned fiber cue.

We further examined the correlation between the traction stress and the speed of cell migration under different conditions (**Figure 7I**). We recorded time-lapse imaging videos for 8 h at each condition and analyzed the average speed and traction stress. Surprisingly, we did not find any noticeable pattern that correlates the traction stress with speed across different cells under various conditions (MM, AF and RF). Nevertheless, the control experiment along with formin and Arp2/3 inhibition in HFF and HaCaT cells on AF exhibited a linear pattern, suggesting that traction stress and speed are correlated when a directed cue is presented (**Figure 7I**). Similar results are shown for directionality, which appears to be linearly correlated to traction stress in each cell type (**Figure 7J**). To further strengthen our results and examine whether this correlation is consistent across different cell lines, MDA-MB-231 cells were plated on the AF surface at 200 Pa, 2000 Pa, and 20000 Pa. Interestingly, we found that both speed and directionality increased with increased traction stress (**Figure 7K&L**), corroborating the behavior seen in HFFs and HaCaT(**Figure 7I&J).** Overall, while traction stress does not seem to correlate with speed and directionality across different structural organizations of collagen and across cell lines, using one cell line and examining one type of structural organization (aligned collagen fibers), traction stress positively correlates with both speed and directionality.

## 3. Discussion

Our work characterizes the traction stress magnitude and kinetics exerted by cells on fiber networks that have been poorly explored to date. We outline the relationship between traction stress, directionality and migration speed across several different organizational patterns of collagen by employing d-TFM. The key advantages of d-TFM over other traditional 2D TFM is that it is amenable to examining traction forces on collagen fiber networks and measures traction force from displacements of beads attached to the ECM on top of the flexible substrate and as well as from displacements of beads within the underlying flexible substrate. Traditional 2D TFM has been a powerful technique to measure cellular traction force since its discovery in 1980 ^59^. However, a limitation of 2D TFM is that it most commonly measures cell traction stress on flexible substrates with adsorbed ECM molecules, lacking the structure of assembled and aligned ECM fibers. The alignment of collagen fibers resulting in contact guidance, has a significant impact on cell traction stress ^19,23^. While some studies have reported the traction stress on anisotropic substrates ^23,25^, no work has yet benn reported on aligned collagen fiber networks. Our d-TFM approach allows for the characterization of forces transmitted through collagen fiber networks, leading to an understanding of how fiber-induced forces stimulate cells to engage in contact guidance in physiological environments.

We first examined the deformations induced by MDA-MB-231 cells on aligned collagen fiber networks on three substrates: mica, glass and 2000 Pa d-TFM PAA gels. The results showed that there are similar deformations exerted by MDA-MB-231 cells on aligned collagen fibers on mica in comparison to those on d-TFM substrates at 2000 Pa. This suggests the cells sense an observed modulus of collagen fibers on mica that is similar to 2000 Pa (**Figure 1M**). No deformations were observed when collagen fibers were transferred to glass (**Figure 1M**). Previously, our lab and others indicated that collagen fibers can be deformed due to stochastic, but sparse interactions between the collagen fibers and the mica surface, mimicking a flexible substrate ^35,60^. Our results are consistent with these reports, confirming the unique characteristic of mica as presenting flexible collagen fibers for the study of cell-induced collagen fiber deformation. However, we were able to quantitatively benchmark deformations of collagen fibers on mica with a flexible substrate of known elastic modulus. Even though mica may not be an effective substrate for highly contractile cells such as HFFs due to the delamination of collagen fibers from mica, the covalent attachment of collagen fibers to flexible substrates blocks the delamination, while presenting similar mechanical properties (**Figure 2E** and **Figure S4A**). Using our d-TFM at 2000 Pa, we demonstrated that the displacement on aligned collagen fibers (BAF) is greater than that on flexible substrates (BEG) (**Figure 1I&M**). The ratio of these displacements is nearly same in two different cell lines (MDA-MB-231 and HFF), even though the cells have different sizes, shapes and contractile forces (**Figure 1I**). The ratios are also similar across three different collagen organizations (**Figure 4N**). This demonstrates that the difference in displacements observed from beads at the ECM layer and those within the flexible substrate is a property of the system and is likely due to the dissipation of stress in the flexible substrate, resulting in different displacements at different depths. As a result, using displacements from depths further into the PAA gel to generate traction stresses may compromise the precise calculation of traction stress at the cell-ECM interface (**Figures 2-4**) ^32^. We demonstrated that each cell exerts spatially heterogeneous traction stresses on aligned fibers (**Figure 3A&B**), with the average traction stress depending on stiffness (**Figures 2G and 3C, D**). This is consistent with what others have shown on adsorbed ECM monomers ^42,43^. Taken together this system has revealed our ability to measure traction stress on aligned fiber networks and to assess the mechanical coupling between the fibers on the top of the substrate with the substrate itself, showing a stiffness-dependent traction stress response that is seen on non-fiber systems.

Fiber networks are much different from isotropically adsorbed monomers in their biophysical characteristics. In addition, aligned fiber networks might elicit very different force transmission characteristics. Therefore, we explored differences between isotropically adsorbed collagen monomers, aligned collagen fibers and randomly organized collagen fibers. There is some indication that traction force exerted from anisotropic substrates exceeds the traction force exerted from isotropic substrates, but no strong evidence is available to validate this claim in fiber networks ^23^. Using myosin phosphorylation as a proxy for traction force, a recent study suggested that a greater traction force is exerted on random fiber networks as compared to aligned fibers but this report did not directly measure traction stress ^25^. By comparing traction stress on isotropically adsorbed collagen, aligned collagen fibers and random collagen fibers, our results establish that higher traction forces are indeed exerted on random collagen fibers (**Figure 4H-J**). Surprisingly, results revealed that traction stress kinetics during spreading are independent of stiffness (**Figures 2G&H**); rather, they depend on how the collagen is presented, whether as monomers, as aligned fibers or as randomly organized fibers: uniform collagen (BMM vs BEG), aligned collagen fibers (BAF vs BEG) or random collagen fibers (BRF vs BEG) (**Figures 4H-J**). Traction stress kinetics on aligned fibers and isotropically adsorbed collagen are approximately similar and both of them were fitted best using an exponential function dependent on spreading time, reaching a steady state at 2 h. Traction stresses on random collagen fibers continue to increase linearly with spreading time. Differences in magnitude and kinetics are likely due to distinct mechanisms involved in cell-ECM fiber interactions in each structural condition. Random collagen fibers result in F-actin stress fibers oriented in multiple directions, due to numerous adhesion contact points. Cells actively pull collagen fibers close to their body after initial cellular adhesion in order to better spread and stabilize adhesion, accumulating many numerous adhesions oriented in different directions ^25,61^. We propose that the highest traction stress is on RF substrates because of multidirectional constrained FAs (MCFAs) transmitting force between the cell and the ECM. In contrast, on aligned collagen fibers, cells tend to elongate and polarize in one direction towards the fiber direction ^62^, transmitting force via directional constrained FAs (DCFAs) thereby stabilizing ECM interactions within 2 h, reaching a steady-state level of traction stress. Multidirectional fiber ECMs collectively transmit signals through MCFAs, generating stronger adhesive strength than directional constrained FAs (DCFAs) on AF and unconstrained FAs (UCFAs) on MM (**Figure 4&8**). In addition to the magnitude and kinetics of traction stress on different substrates, we explore the coupling between traction stress and area. Several studies showed that the traction forces on uniform substrates are often linearly correlated with the cell spreading area ^42,43^, consistent with our results on uniform substrates (**Figure 4O**). However, we discovered that traction forces exerted on fiber networks do not correlate with the cell area (Pearson’s correlation < 0.2) (**Figures 2I, J, and 4O**). This correlation is even weaker when the stiffness is higher (**Figures 2I&J and 8**). This is likely due to the directional bias of the traction stress of aligned cells, resulting in an unequal force distribution along fibers and perpendicular to fibers in contrast to uniform substrates where cells spread and orient freely in all directions. Consequently, collagen fibers and, in particular, aligned fibers appear to have distinct traction stress signatures that are not observed when examining traction stress on isotropically adsorbed collagen.

To understand the role of the cytoskeleton in regulating traction stress formation on aligned collagen fibers, we examined formins and Arp 2/3. Formins and Arp2/3 generate distinct F-actin networks, the former creating linear F-actin networks and the latter creating branched F-actin networks. Our initial hypothesis was that branched and linear F-actin networks compete for a common pool of G-actin, so inhibiting one will result in the building of another. Furthermore, we predicted that Arp2/3 builds branched networks that facilitate an ability to make directional changes, but generate small traction stresses, whereas formins build linear networks that facilitate directional migration by generating large traction stresses. By inhibiting Arp2/3 (CK-666) and formins (SMIFH2, FHOD3 KD) in HFF cells on aligned collagen fibers, we found that Arp2/3 and formin diminish the peak traction stress, but not the average traction stress in a model mesenchymal cell line, HFFs. This indicates that low traction stress areas increase in level and compensate for the reduced high traction stress magnitude, leading to an insignificant change in the average traction stress. These findings likely suggest that the conflicting effects of formins and Arp2/3 tend to balance the force magnitudes, resulting in the cells performing approximately equal amounts of work on average ^43^ (**Figure 5C**). In epithelial cells (HaCaTs), a different story emerges. We showed that these F-actin regulatory proteins significantly affect the kinetics of traction stress as well as the magnitude of traction stress (**Figure 6E, F and 8**). Specifically, inhibiting Arp2/3 slows the increases of traction stress during spreading. As a result, cells reach a steady state slower compared to the control group. Alternatively, inhibiting formins diminishes the magnitude of the traction stress at steady-state. The magnitude of stress is directly controlled by the number of stress fibers, which act as a driver to connect FAs and function as a source of traction force generation, anchoring to the substrate ^63–65^. Interestingly, inhibiting formins not only increases the traction stress magnitude, but increases the correlation between traction stress and cell area (**Figure 6I&J and 8**). Likely, diminished formin activity decreases the ability of cells to elongate, so spreading proceeds as it might on isotropic substrates, which do show a strong correlation between cell area and traction stress. In addition to changes in traction stress kinetics, steady-state magnitude and traction stress-area coupling, formins and Arp2/3 alter the cells ability to move between parallel alignment with collagen fibers and perpendicular alignment to collagen fibers. Cells transmit higher traction stress when transitioning between parallel and perpendicular alignment (**Figure 6D&E**). During this transitional state, the cell has to perform three steps: relaxation of traction stress, protrusion turning and acceleration of movement. Cells may increase traction stress to generate the necessary force needed to overcome the bending or pivoting of linear F-actin fibers. During turning, parallel fibers act as a bridge where the cell body is likely to hang at the bridge edge. Lack of adhesive support facilitated by strong traction stress results in inward contraction and consequently activated stronger intracellular force ^25,66^. Interestingly, blocking both of these F-actin regulatory proteins decreases the turning stress, which demonstrates their prominent role in cell turning from parallel to perpendicular and vice versa (**Figure 6I**). Taken together formins and Arp2/3 play distinct roles in mesenchymal and epithelial contact guidance. In mesenchymal cells formins and Arp2/3 play roughly overlapping roles with respect to traction force generation, whereas in epithelial cells they at times have overlapping roles as in controlling traction stress during cell turning, but they also have distinct roles in governing traction force kinetics and steady-state magnitude.

Finally, our work establishes the relationship traction stress and directionality of cells, as well as between traction stress and speed on aligned collagen fibers. Several studies have shown that the efficiency of contact guidance in directing cell alignment depends on factors such as the architecture of cues (micro-patterned, grooves/ridges, and aligned fibers), cell type and intracellular actomyosin activity ^23,27,35,39,47,67,68^. It is known that contact guidance can be enhanced by upregulating both adhesion and contraction regulated by the F-actin cytoskeleton ^27,54^, even if the initial steps of contact guidance are independent of contractility ^69^. While adhesion and contraction are intimately linked to traction force generation, the direct link between traction stress generation and contact guidance is not known. We determined that aligned fiber-driven traction stress controls the directionality index of cells, by investigating three different structural cues: uniform collagen monomers (MM), aligned collagen fibers (AF), and random collagen fibers (RF). Greater directionality was observed on AF. The lack of aligned fibers on MM and RF substrates leads to marginal directionality on these surfaces. It appears that a threshold level of traction stress is required for directional migration, but high traction stress is not sufficient, there must be aligned collagen fibers present in order to direct migration (**Figure 7H**). Traction stress was also shown to increase directionality index when examining cells on different substrate stiffnesses and across cytoskeletal perturbation (**Figure 7J&L and 8**). In addition to directionality, traction stress also affects the migration speed on aligned collagen fibers. A myriad of studies report a biphasic relationship between migration speed and stiffness across different cell lines ^15,16,19^. We investigated this correlation to see if this was true on aligned fibers. We did not observe a biphasic relationship between traction stress and migration speed (**Figure 7I**). Pharmacological inhibition of formin and Arp2/3 in HFF and HaCaT cells on d-TFM at 2000 Pa, as well as in control MDA-MB-231 cells results at d-TFM at 200 Pa, 2000 Pa, and 20000 Pa on AF surfaces, collectively revealed a linear correlation between traction stress and migration speed on aligned collagen fibers. Cell migration speed increased as traction stress increased within examined elastic modulus (**Figure 7I&K and 8**). Overall, our results show that increases in traction stress increase both cell directionality and speed. Additionally, a minimum level of traction stress is needed to drive marginal fidelity directional cell migration on aligned collagen fibers, but increasing from marginal to high levels of directional migration require no further increases in traction stress.

## 5. Conclusions

In summary, we developed a d-TFM system and measured the traction stress exerted on aligned collagen fiber networks. We revealed that traction stress exerted on both aligned and random collagen fiber networks was greater than those exerted on isotropically adsorbed collagen. Furthermore, traction stress measured at the cell-ECM interface was much larger than that measured within the flexible substrate, indicating the underestimation of previously measured traction stress. We discovered that while traction stresses on aligned collagen fibers are stiffness-dependent as seen on isotropically adsorbed collagen, the area does not correlate with traction stress magnitude. Formins and Arp2/3 share overlapping roles in regulating traction stress in mesenchymal cells and during epithelial cell turning, but have distinct roles in affecting the kinetics and magnitude of traction stresses. Finally, we established a correlation between traction stress, cell directionality, and cell speed on aligned collagen fibers. Traction stress is positively correlated with the directionality of cells on aligned collagen fibers, but only at low levels of directionality. Speed is also positively correlated with traction stress, but does not show biphasic behavior, unlike isotropically adsorbed collagen. Taken together, these findings reveal distinct traction stress signatures that are present when collagen fibers are aligned.

## Author Contributions

GN: Conceived experiments, conducted experiments, analyzed and interpreted the data across all figures, prepared the figures, wrote and edited the article. AF: Wrote code, conducted AFM experiments and assisted in analyzing data. FN: Performed the FHOD3 knockdown experiment, analyzed and interpreted the data. I.C.S.: Conceived experiments, analyzed and interpreted the data, acquired the funding and edited the article.

## Acknowledgements

GN and ICS thankful to the National Institutes of Health (Grant No. GM143302) for supporting this work. We acknowledged all our lab members for the discussion and valuable feedback.

## Abbreviations

ECM: Extracellular Matrix
DPBS: Phosphate-Buffered Saline
BME: (β-MercaptoEthanol)
APS: Ammonium Persulfate
APTES: 3-AminoPropylTriEthoxySilane
EGTA: Ethylene Glycol bis(2-aminoethyl ether)-N,N,N′,N′-Tetraacetic Acid
PVDF: Polyvinylidene DiFluoride
BSA: Bovine Serum Albumin
TEMED: N,N,N’, N’-Tetraacetylethylenediamines
TBS: Tris Buffered Saline
HEPES: 2-hydroxyethylpiperazine-1-ethane sulfonic acid
FBS: Fetal Bovine Serum
DMSO: DiMethyl SulfOxide
MES: 2-N-Morpholino Ethane Sulfonic acid
ECL: Enhanced ChemiLuminescent
HFF: Human Foreskin Fibroblast
EMT: Epithelial to Mesenchymal Transition
AFM: Atomic Force Microscopy
PAA: PolyAcrylAmide
MM: Monomers
AF: Aligned Fibers
RF: Random Fibers
TFM: Traction Force Microscopy
d-TFM: dual Traction Force Microscopy
e-blebb: Blebbistatin

