## Supplemental Methods and Figures for "Aligned collagen fibers drive distinct traction force signatures to regulate contact guidance"

### 1. Materials and Methods:

##### 1.1 Assembling collagen on mica

A freshly cleaved 15 mm x 15 mm piece of muscovite mica (highest grade VI, Ted Pella, Redding, CA) was used for the epitaxial growth of collagen fibers. Rat tail collagen type I (Corning, Corning, New York) was diluted using 50 mM Tris-HCl (Fisher Scientific, Waltham, Massachusetts) and 200 mM KCl (Fisher Scientific, Waltham, Massachusetts) to achieve final concentrations of 10 µg $ml^{-1}$ (for aligned collagen fibers) or 100 µg $ml^{-1}$ (for random collagen fibers) and added to the top of the mica. Following the 12-h incubation period, the mica was washed with nano-pure water. The mica was then placed inside a tissue culture dish, allowed to dry overnight and was ready for either transfer to other substrates or for attaching of fluorescence beads.

##### 1.2 Transferring collagen fibers from mica to functionalized glass

First, 22 mm × 22 mm coverslips (Corning, Corning, New York) were treated with piranha solution (3:1 $H_2SO_4$ (Fisher Chemical) to $H_2O_2$ (Fisher Chemical)) for 1 h to ensure thorough cleaning and oxidation. After treatment, the coverslips were immersed in 1% (v/v) 3-aminopropyltriethoxysilane (APTES, Thermo scientific) solution prepared in 1 mM acetic acid (Alfa Aesarfor, Ward Hill, Massachusetts) for 2 h to facilitate silanization. The coverslips were then thoroughly rinsed with deionized water and allowed to air dry. Subsequently, the coverslips were baked in an oven at 100 °C for 1 h to promote covalent bonding of the APTES layer. Once cooled, the coverslips were treated with 6% (v/v) glutaraldehyde (Electron Microscope Science, Hatfield, Pennsylvania) in Dulbecco's Phosphate-Buffered Saline (DPBS) for 2 h to introduce aldehyde functional groups. Following this treatment, the coverslips were washed with deionized water to remove residual glutaraldehyde. Coverslips were good for two weeks. For transferring the collagen, 30% gelatin (Sigma Aldrich, St. Louis, Missouri) was dissolved in water and heated to 40 °C for 1 h, and 500 µl of the gelatin solution was placed on the mica with assembled collagen. After a 1-h incubation, the gelatin was peeled off and placed on the chemically functionalized glass. Following an additional 1 h of incubation of the gelatin on the target substrate, the samples were washed with 3 ml of DPBS lacking magnesium and calcium (Gibco, Waltham, Massachusetts) by placing the DPBS inside the culture dish containing the substrate and incubating at 37 °C for 2 h. After this period, the underside of the sample was cleaned with a Kim wipe (Kimberly-Clark, Irving, Texas). The samples were further washed for an additional 45 min, exchanging the DPBS at intervals of 15 min to remove gelatin from the surface as much as possible. It is crucial to eliminate all the gelatin, otherwise, fluorescence beads in the next step will attach to gelatin instead of collagen fibers, resulting in their easy removal when cells and media are added. The samples were then ready for attaching fluorescence beads.

##### 1.3 Transferring collagen fibers from mica to functionalized polyacrylamide gels

First, 22 x 22 mm coverslips were covered with 0.1 M NaOH for 10 min. Following this, the coverslips were treated with a 5% APTES solution for 5 min. The coverslips were then washed three times with deionized water, each wash lasting 5 min. Next, a 0.5% glutaraldehyde solution was applied to the coverslips for 30 min, followed by additional washes with deionized water. The solution of stock beads (200 nm FluoSphere™-dark red 660/680, 2674015, Invitrogen- Thermo Fisher Scientific) was sonicated for 5 min followed by 1 min of vortexing. This we done three times. To prepare beads embedded in polyacrylamide (PAA) gels of different elastic moduli, fluorescence beads (200 nm) were diluted 1:50 in the PAA solutions. PAA solutions of 3%, 5% and 8% acrylamide and 0.03%, 0.1%, and 0.26% bis-acrylamide were prepared with 0.05% ammonium persulfate (APS, Bio-Rad, Hercules, California) and 0.15% N,N,N', N'-Tetramethylethylenediamine (TEMED, Fisher Scientific), respectively, to achieve gels with elastic moduli of 200, 2000, and 20000 Pa [38]. The functionalized coverslips were placed face down onto drops of these solutions on microscope slides, allowing polymerization to occur for 30 min. The samples were then treated one time with 2 mM sulfo-SANPAH (Thermo Scientific) under UV light for 15 min to functionalize their surfaces. Collagen fibers were transferred as described above for glass or coverslips were exposed to 100 µg $ml^{-1}$ collagen dissolved in 0.01 M acetic acid overnight in the fridge. The samples were then ready to attach fluorescence beads.

**1.4 Attaching fluorescence beads to collagen monomers or fibers on mica, glass or PAA gels**

The solution of stock beads (40 nm FluoSphere™-red 580/605, 2752655, Invitrogen-Thermo Fisher Scientific) was prepared in a similar way to the 200 nm beads above. A 1:5000 dilution of stock bead solution was dissolved in nanopure water and added to the substrates. Collagen fibers or monomers on mica, glass or PAA gels were placed on 10 mm petri-dish and 2 ml of bead solution was added on the top of the substrate and incubated at room temperature for 4 h or overnight at 4 $^{o}$C. Following the incubation period, the substrates were washed with nanopure water at least three times and the substrate was ready for cell seeding. Note that the concentration of embedded beads and bead solutions for adsorption onto fibers was tuned to achieve a similar density of ~1 bead/$\mu m^2$.

**1.5 Bead density analysis**

Bead density was quantified by dividing the number of beads detected in a field of view by the size of the field of view. The number of beads for each field of view was measured in Fiji using the "find maxima" option. The threshold used was tuned for each data set so that only beads were counted.

**1.6 Characterizing collagen fibers using Atomic Force Microscopy (AFM) and immunofluorescence**

The topography of the collagen fibers on mica and transferred to glass was imaged using an AFM (Digital Instruments, now Bruker Nano, Santa Barbara, CA) multi-mode AFM in tapping-mode with TESPA probes with 40 mV drive amplitude and frequency of 320 kHz. Captured images were processed with the plain fit and flattened routines as appropriate. Collagen fibers were also stained and imaged using immunofluorescence. Collagen fiber samples were first blocked with blocking buffer (0.02 g $ml^{-1}$ BSA prepared in 10% Tris-buffered saline (TBS) containing 0.01% Tween-20) for 20 min and then incubated with 1:100 rabbit anti-rat collagen primary antibody (2150-1908, BioRad) for 24 h. After washing three times with 1x TBS for 5 min each, samples were incubated with a 1:200 dilution of donkey anti-rabbit alexa 555 secondary antibody (Invitrogen™ A-31572, Fisher Scientific) for 1 h, followed by additional washes.

**1.7 Culturing cells and treating with inhibitors**

An immortalized human foreskin fibroblast cell line (HFF, ATCC, Manassas, VA), a human mammary carcinoma cell line (MDA-MB-231, ATCC) and an immortalized human keratinocyte cell line (HaCaT, kind gift from Dr. Torsten Wittmann) were used in this study. MDA-MB-231s and HaCaTs were cultured in Dulbecco's modified Eagles medium (DMEM, Sigma Aldrich). It was supplemented with 10% fetal bovine serum (FBS) (Gibco, Grand Island, New York, USA), 1% penicillin–streptomycin (Gibco), and 1% Glutamax (Gibco) at 37 °C in 5% $CO_2$. The same media with 15% FBS (Gibco) was used for culturing HFFs. When live cells were imaged, cells were placed in culture media that lacked phenol red, but contained 15 mM 4-(2-hydroxyethyl)-1-piperazineethanesulfonic acid (HEPES), Glutamax, penicillin/streptomycin, 10% Fetal bovine serum (FBS). This is referred to as clear imaging media. Cells were centrifuged, counted and suspended in culture media 20,000 # $ml^{-1}$ (HFFs) or 30, 000 # $ml^{-1}$ (MDA-MB-231 and HaCaT). SMIFH2 (Millipore), CK-666 (Millipore) and blebbistatin (Millipore) were dissolved at 50 mM, 50 mM and 20 mM stock solutions in dimethyl sulfoxide (DMSO). Stock solutions of the inhibitor were diluted in either culture media or imaging media by at least 1000-fold to make working solutions used on cells.

**1.8 Small interfering RNA (siRNA) transfection**

We used siRNA interference, Silencer® predesigned FHOD3 siRNA oligonucleotides (AM16708, targeting FHOD3, Thermofisher Scientific). FHOD3 siRNA sense sequence: 5′-GGAAGUAGCAGAACCACUCtt-3′ and antisense sequence: 5′GAGUGGUUCUGCUACUUCCtt-3′. According to the manufacturer's protocol, oligonucleotides were transfected in cells using Lipofectamine RNAiMAX (13778075, Invitrogen). Cells were seeded in a 60 mm culture dish using 2 x $10^6$ cells/5 ml and incubated for 48 h. The complex formed by mixing Lipofectamine RNAiMAX and siRNA was added to the cells at an siRNA of 50 μM in the media. The control group involved cells transfected with no siRNA and no Lipofectamine RNAiMAX reagent. Cells were incubated for 48 h with the transfection mix, washed with DPBS and processed for several assays.

**1.9 Total RNA extraction**

Cells were cultured to a confluence of 80%, washed with warm DPBS and trypsinized using 0.25% trypsin-EDTA (Gibco Life Technologies). For adherent cells, $2x10^6$ cells were used to extract RNA using an RNeasy Mini kit (74104, Qiagen, Valencia, CA). Cells were centrifuged and pelleted at approximately 8000 x *g* for 5 min in the centrifuge at room temperature. The cell pellet was lysed by adding Qiagen buffer RLT, followed by equal volumes of 70% ethanol and mixed by pipetting gently. The mixture was transferred to a spin column tube and centrifuged several times for different times using different washing buffers as prescribed by the manufacturer. Finally, the column was put into a new collecting tube and RNA sample was eluted with 30 μL of RNase-free water at 8000 x *g*. The concentration of the collected RNA was quantified using a Nanodrop.

**1.10 Reverse Transcription PCR (RT-PCR)**

For RT-PCR, 2.5 μg of cDNA using SuperScript™ III First-Strand Synthesis System (18080051, Invitrogen) was used. For each reaction, 2.5 μg of RNA, with a final concentration of 1 mM of dNTP, 5 ng $\mu L^{-1}$ of random hexamers were added in the final volume of 30 μL diethylpyrocarbonate treated water. The mix was incubated for 5 min at 65 $^oC$ and 5 min on ice. Following these incubations, 2 x RT buffer,10 mM $MgCl_2$, 0.02 M DTT and 4 μg $\mu L^{-1}$ RNase OUT and superscript III was uniformly mixed and incubated for 50 min at 50 $^oC$, followed by 85 $^oC$ for 5 min. Then, 2 μl of RNase H was added to degrade any remnant RNA and incubated at room temperature for 20 min. To prepare the PCR reaction we used Taq buffer, 0.2 mM dNTP mix, 1.5 mM $MgCl_2$, 5 U/μL Taq polymerase and a 2.5 μg $\mu L^{-1}$ DNA template followed by 3 μM for each primer. The PCR reaction program included the following conditions: Initial denaturing temperature was 94 $^oC$ for 2 min, denaturing temperature was 94 $^oC$ for 1 min, annealing temperature was 55 $^oC$ for 30 min, extension temperature was 72 $^oC$ for 10 min and 30 cycles were used. Gel electrophoresis was used to validate formin knockdown. The samples were run on the 60 mL prepared and loaded agarose gel run at 3.4 V/cm (70 V) for 90 min. Lastly, the gel was imaged by a gel imager (smartDoc, Accuris) and data was analyzed using ImageJ.

**1.11 Immunofluorescence imaging and analysis**

Cells were plated onto target substrates at a concentration of 40,000 # $ml^{-1}$ (HFFs) for subsequent experiments. Cells were fixed and stained to visualize their F-actin structure and to quantify their response to contact guidance through morphology and alignment behavior. First, the cells were treated with 4% paraformaldehyde (Fisher Scientific) for 10 min, prepared by diluting 16% paraformaldehyde in a cytoskeleton buffer consisting of 10 mM 2-N-morpholino ethane sulfonic acid (MES, pH 6.1, Fisher Scientific), 3 mM $MgCl_2$ (Fisher Scientific), 138 mM KCl (Fisher Scientific) and 2 mM EGTA (Sigma

Aldrich). Next, cells were permeabilized using a 0.5% Triton-X (Fisher Bioreagents) solution in the cytoskeletal buffer for 5 min. Unreacted aldehydes were then blocked with 100 mM glycine (Fisher Scientific) for 15 min. Cells were washed three times with TBS for 5 min each. Cells were stained with Alexa 488-phalloidin (Thermo Fisher Scientific) for 1 h and DAPI (Sigma) for 15-30 min in TBS containing 0.1% (v/v) Tween-20 (Fisher Bioreagent) and 2% (w/v) Bovine Serum Albumin (BSA, Sigma Life Science). For anti-paxillin staining, after initial phalloidin staining and washing with TBS, cells were incubated with 1:200 dilution of mouse anti-human paxillin primary antibody (610051, BD Biosciences) for 24 h at 4 °C. Samples were then washed with TBS, followed by incubation with a 1:200 dilution of Cy™5 AffiniPure® donkey anti-mouse IgG secondary antibody (715-175-150, Jackson ImmunoResearch) and 1:500 dilution of DAPI for 1 h. Samples were washed again with TBS three times for 5 min each. To improve image resolution, Prolong Gold mounting media (Thermo Fisher) was applied between the substrate with cells and the cover slides. Samples were sealed with VALAP (a mixture of Vaseline, lanolin, and paraffin) and imaged the next day using epifluorescence microscopy with 10x and 40x objectives ($NA$ = 0.3 and $NA$ = 1.3, Nikon). The immunofluorescence images were analyzed with ImageJ. Lines around the cell's edge were drawn to find the major and minor axis of cell shape. The aspect ratio was calculated as the ratio of the major axis of the cell divided by the minor axis of the cell. The angle of the cell was calculated as the angle between the long axis of the particular cell and average angle of all cells within that window. The Directionality Index (DI) was calculated as the $\cos(2|\Delta\theta|)$, where $|\Delta\theta|$ represents the angular deviation of the cell from the average cell angle in the region. Cells with an aspect ratio less than 1.5 were considered as non-aligned and assigned their DI as zero.

**1.12 Live cell migration imaging and analysis**

Chambers were fabricated by cutting adhesive heat resistant, thin transparent silicone rubber sheets using a craft cutter to form a chamber on a glass surface. This chamber was filled with clear imaging media, supplemented with 10-15% FBS, depending on the cell line and the substrate with the cells was flipped over onto the chamber. The chamber was sealed with VALAP. The setup was then imaged over an 8-h period with images captured at 6-min intervals. This allows for detailed observation and analysis of cell migration behavior. The DI calculations for live cell migration were based on the average of $\cos(2|\Delta\theta|)$, with $|\Delta\theta|$ being the angle difference between the fibers and the cell's direction, derived from changes in $x$ and $y$ positions at 6-min intervals and averaged over 8 h. Speed was calculated by averaging the displacements measured every 6 min over 8 h via Matlab.

**1.13 Displacement and traction stress measurements**

Since the thickness of the fibers was very small (< 5 nm), we considered the elastic modulus of PAA gels with collagen fibers to be the same as the elastic modulus of PAA gels without collagen fibers. This seems reasonable, given collagen fibers have a relatively high elastic modulus, but their bending modulus is much lower. Since the collagen fibers are attached to the flexible PAA gel, their bending and deformation will transfer force to the underlying PAA gel. This means the effective elastic modulus experienced by the cells when on the collagen fibers is essentially the elastic modulus of the underlying PAA gel. Consequently, the role of collagen fibers is to guide cells and induce an anisotropic environment, but not to increase the elastic modulus.

We used Fiji/ImageJ to track the bead displacements and measure the traction stress. Bead tracking within the search window was done using spatial correlation tracking [31]. Initially, images of fluorescent beads in stressed and unstressed gel conditions were stacked. We recorded video from early adhesion, right after the cell adhered to the surface. Consequently, the first image represented the unstressed state of the gel, while the subsequent images captured the stressed state. The stacked images were aligned using Scale-Invariant Feature Transform (SIFT) available under the registration plugin in ImageJ. Using the particle image velocity (PIV) and Fourier transform traction cytometry (FTTC) plugins in Fiji, strain fields and traction stress maps were generated [32]. Briefly, the first staked image was processed using the PIV plugin with the correlation coefficient iteration option (interrogation window sizes: 128 pixels first round, 64 pixels second round, and 32 pixels third round). After that, the produced PIV text file was saved and plotted using the plot function as a displacement map. The normalized median test option was used (parameters used: 0.2 for noise and 5.0 for threshold). The ImageJ FTTC plugin with parameters of Poisson ratio = 0.5; elastic modulus = 200 Pa, 2000 Pa, and 20000 Pa and regularization parameter = $4.0 \times 10^{-10}$ was used to generate the traction force maps. By selecting a region of interest (typically traction stress within the cell region), we measured the average and maximum stress using FTTC.

**1.14 Analysis of traction stress as a function of cell angle with respect to the collagen fiber angle**

Different cells aligned with different angles on aligned collagen fibers at the same time. We categorized the cell orientation angle in 10 discrete angular intervals, e.g. $0^0 < \theta < 10^0 = 5^0$ and $81^0 < \theta < 90^0 = 85^0$. We counted the number of cells within a defined range of angles and calculated the average traction stress to plot against that angle. The cell population distribution against the angle was plotted as the total number of cells within the defined range of angles.

**1.15 Statistical methods**

Statistical significance was assessed using a 95% confidence interval, calculated from the experimental data and placed on graphs in the form of error bars. $p$-values were evaluated by using a two-tailed unpaired student t-test. $p < 0.05$ represents statistically significant difference and $p > 0.05$ represents statistically insignificant. The $p$-value for the difference between two correlation coefficients was computed as follows.

First, we convert Pearson's correlation coefficients ($r_1$ and $r_2$) to Fisher's $Z$ values using the following equation:

$$Z = 2\ln\frac{1+r}{1-r}. \tag{1}$$

Next, the standard error ($SE$) for Fisher's $Z$ values is

$$SE = \sqrt{\frac{1}{n_1 - 1} + \frac{1}{n_2 - 1}}, \tag{2}$$

where $n_1$ and $n_2$ are the sample sizes for the two correlation coefficients. The $Z$-score was for the difference between two Fisher's $Z$ values is

$$Z = \frac{Z_1 - Z_2}{SE}. \tag{3}$$

Next, the $p$-value is obtained by looking up the Z-score in a standard normal distribution table using the following equation:

$$p = 2\left(1 - \phi\left(Z\right)\right), \tag{4}$$

where $\Phi(Z)$ can be found from a normal distribution table.

The p-value for the difference between two kinetic rate constants was computed as follows. The $t$-statistic was found using the following equation:

$$t = \frac{k_i - k_2}{\sqrt{\frac{SD_1^{\,2}}{n_1} + \frac{SD_2^{\,2}}{n_2}}}, \tag{5}$$

where, $k_1$ and $k_2$ are the kinetic rate constants, $SD_1$ and $SD_2$ are the standard deviations computed from the fit and $n_1$ and $n_2$ are sample size. The degree of freedom ($df$) for an independent t-test can be approximated as,

$$df = n_1 + n_2 - 2. \tag{6}$$

The $p$-value was computed in excel using the $t$-statistic, the difference in values ($k_1$-$k_2$) and $df$.

#### 2. Supplementary Figures:

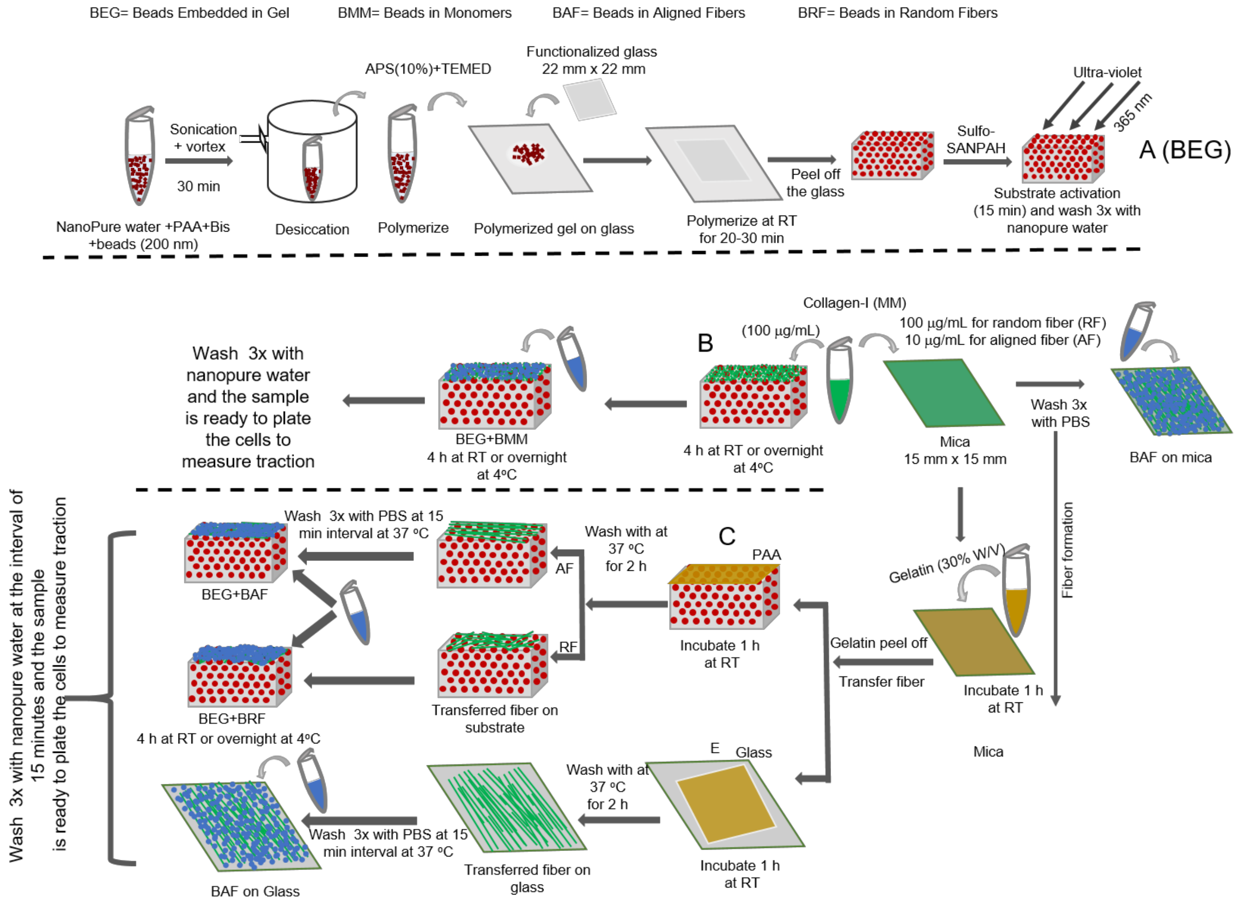


**Figure S1: Schematic showing the overall method creating dual-traction force microscopy (d-TFM) substrates**. (**A**) Embedding beads in PAA gels (BEG) and activating them for either collagen fiber transfer or collagen deposition (MM). (**B**) Adsorption of beads on aligned collagen fibers on mica (BAF on mica) or collagen monomers attached to functionalized PAA gels with embedded beads (BEG and BMM). (**C**) Aligned fibers can either be transferred to functionalized PAA gels with embedded beads (BEG and BAF or BEG and BRF) with subsequent bead adsorption or can be transferred to functionalized glass with subsequent bead adsorption (BAF on glass).

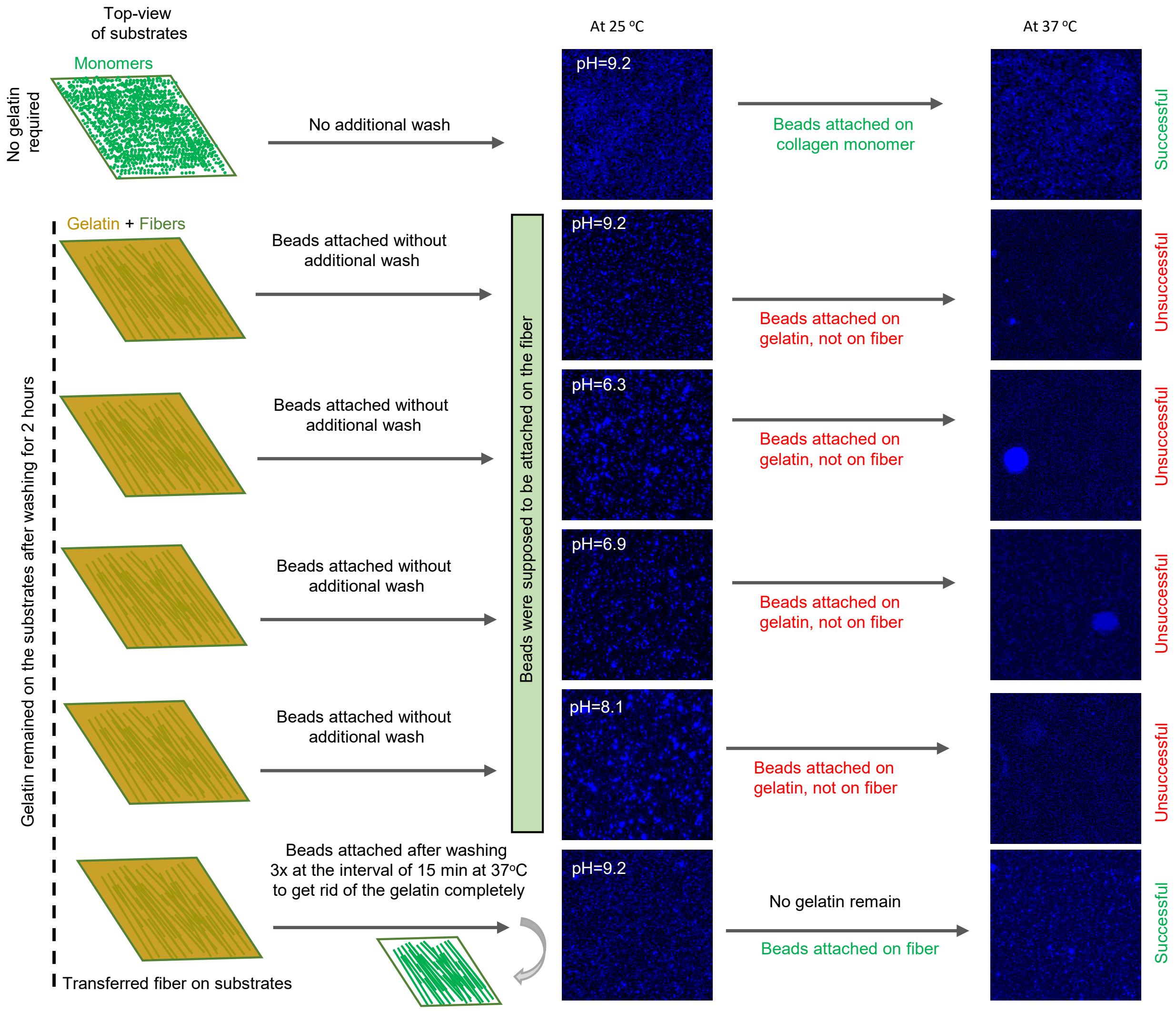


**Figure S2: Schematic figure of attaching fluorescent beads on collagen fiber networks.** While the beads were easily adsorbed to the aligned collagen fibers on mica, attaching them to collagen fibers transferred to glass and flexible substrates was a challenging task due to unwashed gelatin that remained on the surface, even after several washing steps. As a result, beads adsorbed to the gelatin rather than fibers and desorbed once the gelatin was melted away at 37 °C. Thus, additional care was taken while attaching beads to collagen fibers on flexible substrates. An additional step beyond bead adsorption on mica involved performing a wash-and-incubate procedure at 37 °C three times, with 15-minute intervals, to ensure complete removal of gelatin from the fibers attached on glass and flexible substrates.

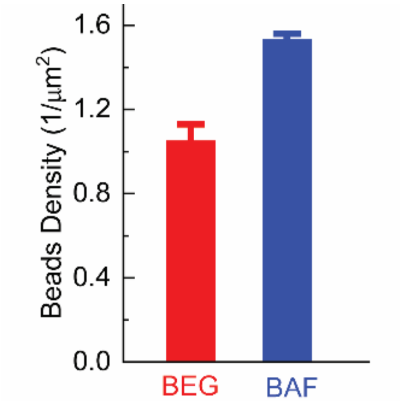


**Figure S3: Bead densities inside flexible substrates and on collagen fiber networks on d-TFM 2000 Pa.** Error bars represent 95% confidence interval. $N_{sample}$= 1, $N_{images} \geq 10$.

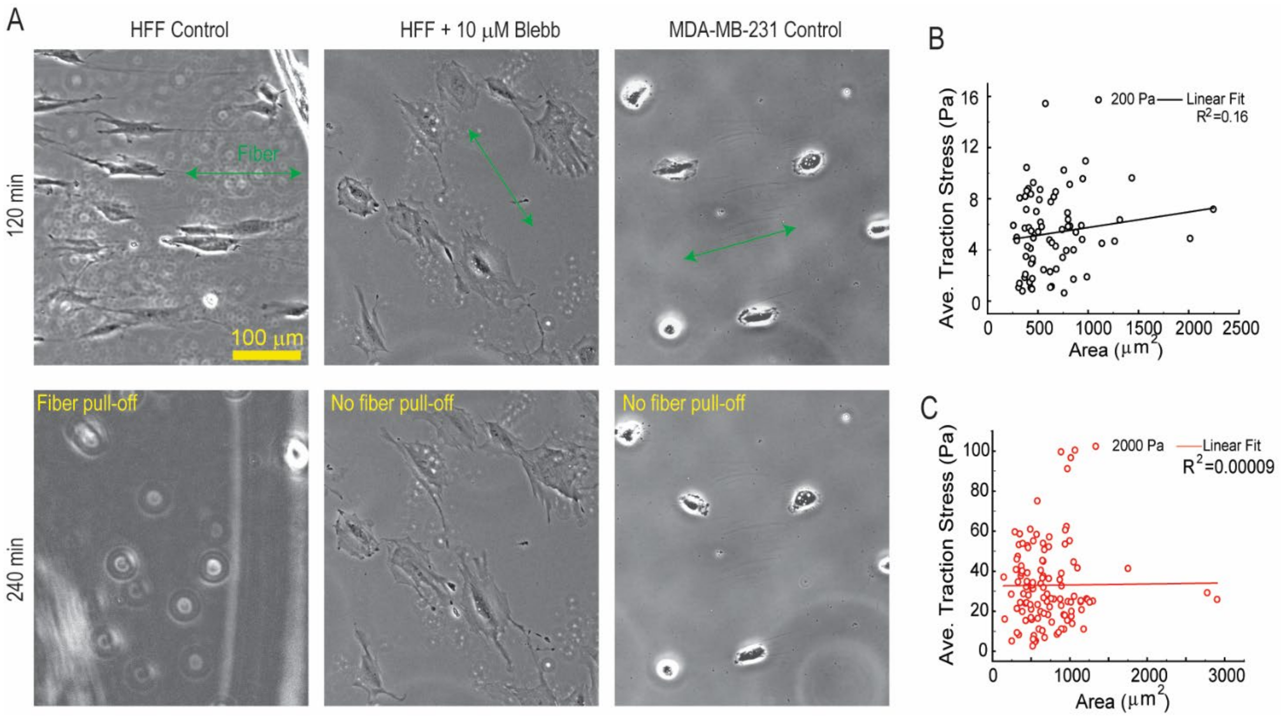


**Figure S4: Inhibiting myosin II limits delamination of collagen fibers by HFFs.** (**A**) HFF and MDA-MB-231 on collagen fibers assembled on mica. (**B**, **C**) Correlation between average traction stress of MDA-MB-231s and area on aligned collagen fibers transferred to d-TFM 200 Pa and d-TFM 2000 Pa.

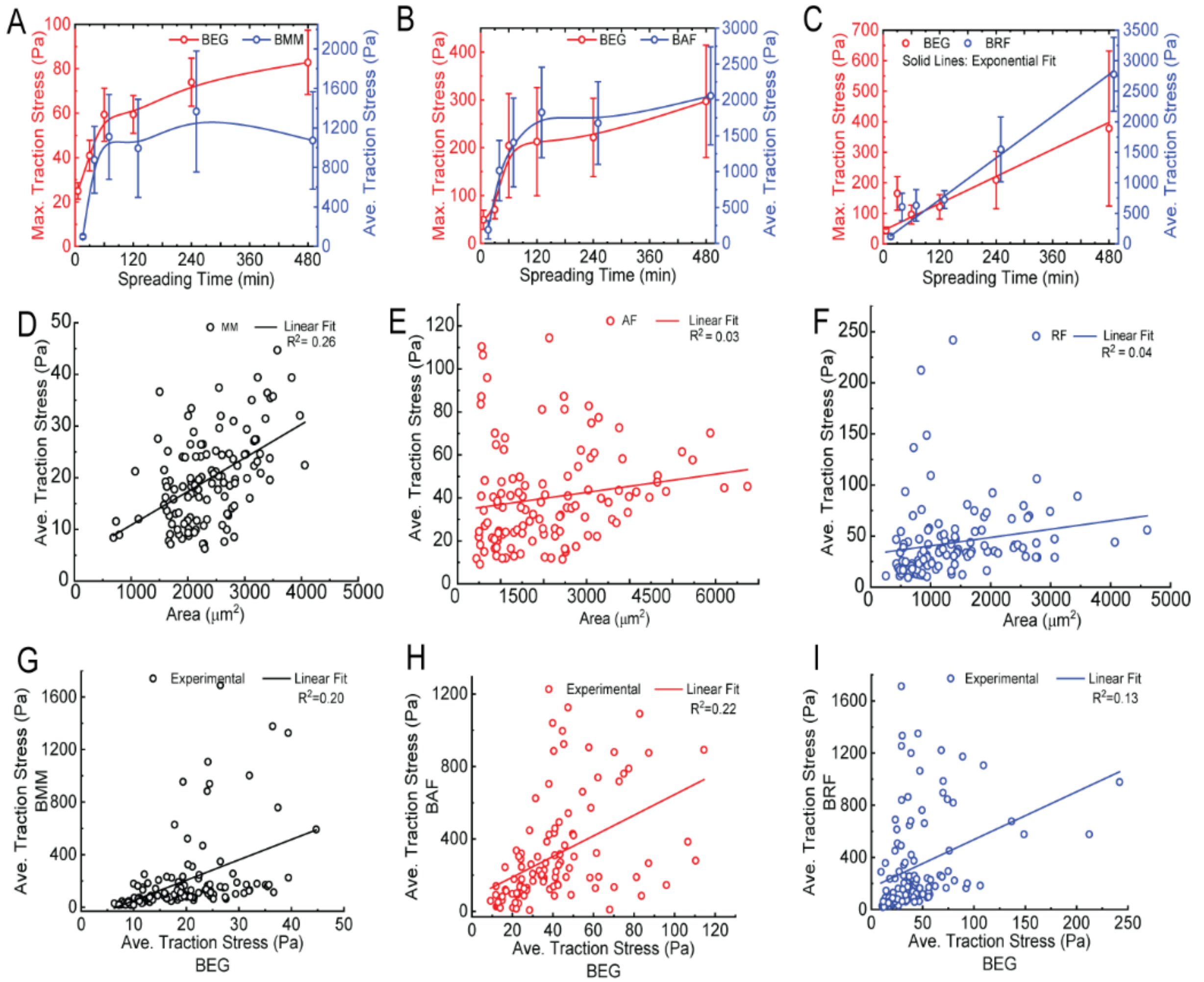


**Figure S5: Traction stress kinetics of HFF cells in response to different collagen organizational structures.** (**A-C**) Maximum traction stress kinetics over time in different collagen organizational structures: MM, AF, and RF. (**D-F**) Correlation between traction stress (BEG) and cell area on MM, AF, and RF at d-TFM 2000 Pa, and (**G-I**) correlation between traction stress exerted on BEG and BAF surfaces on different collagen organizational structures: MM, AF, and RF.

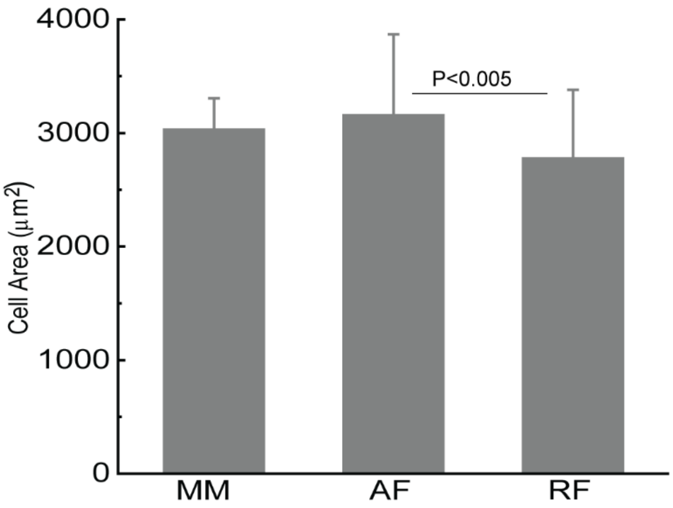


**Figure S6: HFF cell area on different collagen organizational structures:** Cells area on different collagen organizational structures: MM, AF and RF at 480 min on d-TFM 2000 Pa. Error bars represent 95% confidence interval. $N_{sample}$= 3, $N_{images} \geq 20$.

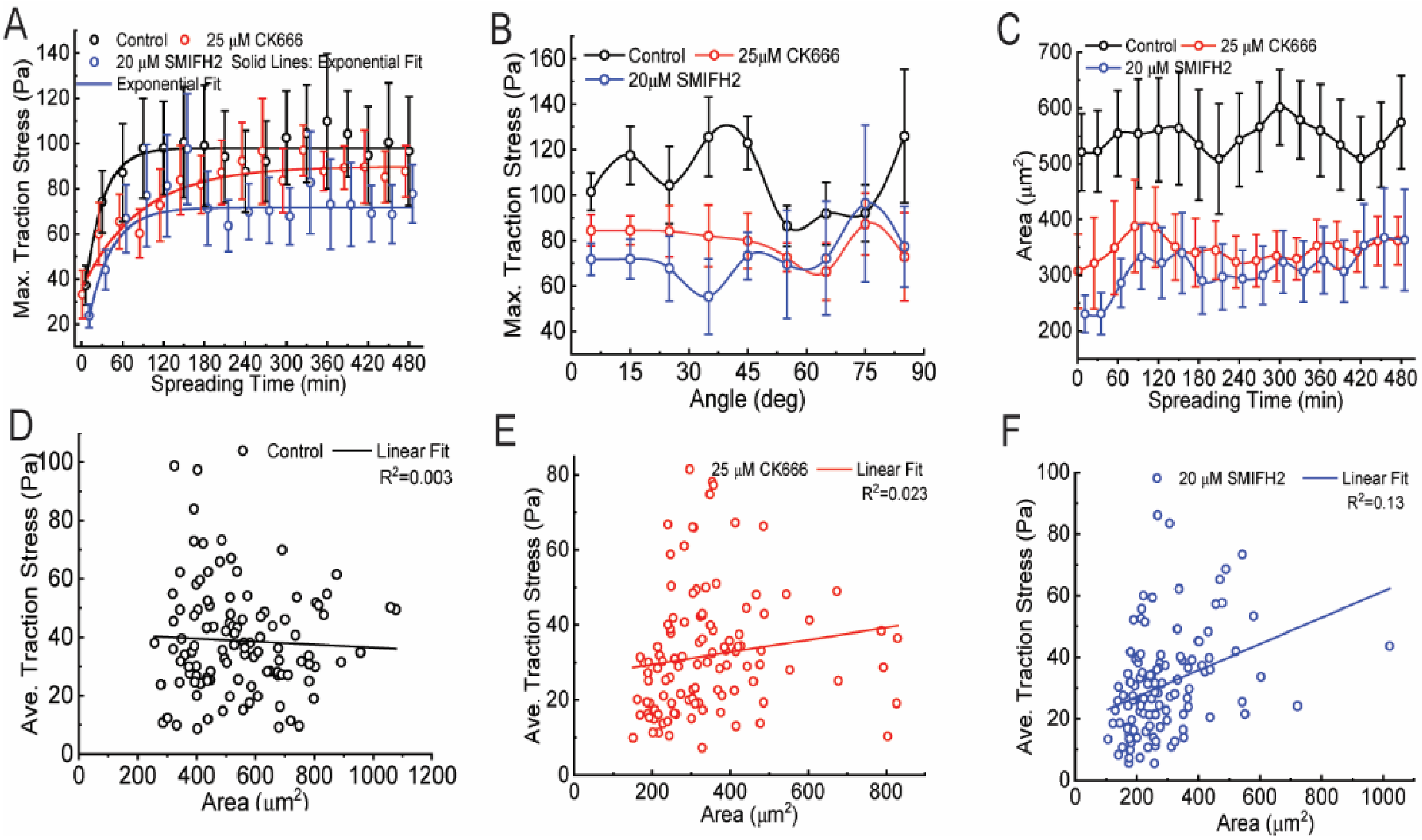


**Figure S7: HaCaT traction stress on aligned collagen fibers in response to F-actin polymerization inhibitors.** (**A**) Maximum traction stress (BEG) kinetics over time on AF at 2000 Pa. (**B**) Maximum traction stress exerted at different angles on AF at 2000 Pa. (**C**) Cell area kinetics over time on AF on d-TFM 2000 Pa, and (**D-F**) correlation between distribution of traction stress and cell area on AF on d-TFM 2000 Pa. Error bars represent 95% confidence interval.

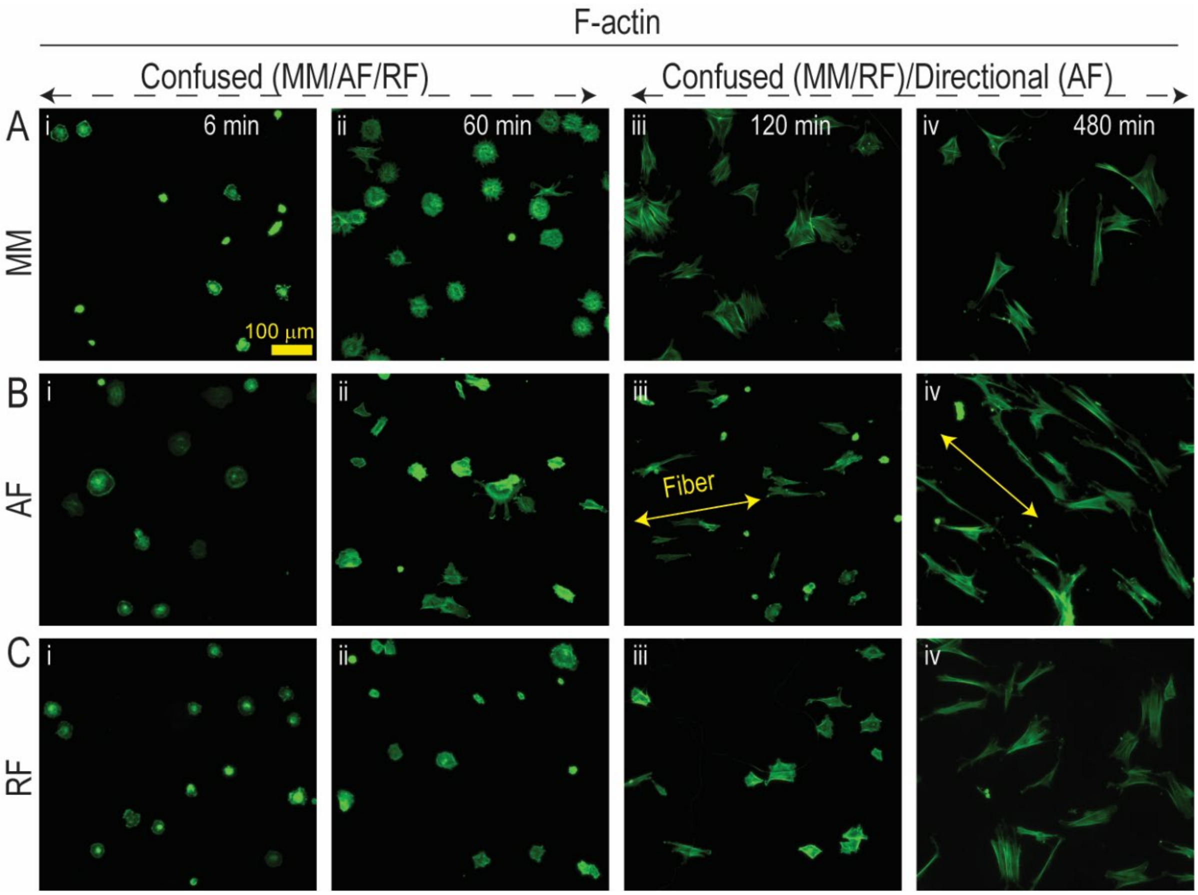


**Figure S8: F-actin organization in HFF cells.** F-actin staining of HFF cells on different collagen organizational structures: MM, AF, and RF at different times. Cells were fixed and stained with phalloidin at different spreading times on d-TFM 2000 Pa.
